# Efficient targeting of heart lesions with cardiac myofibroblasts: Combined gene and cell therapy enhanced by magnetic steering

**DOI:** 10.1101/2024.02.16.580672

**Authors:** M. Schiffer, K. Wagner, E. Carls, J. Nicke, M. Hesse, R. Fratila, S. Hildebrand, A. Pfeifer, D. Eberbeck, M. Malek Mohammadi, J.M. De la Fuente, B.K. Fleischmann, W. Roell

## Abstract

The cardiac scar is a collagen-rich area, which is populated by myofibroblasts and has proven little amenable for therapeutic interventions. Herein, we have established an efficient targeting strategy for cardiac lesions by genetically manipulating embryonic cardiac myofibroblasts (mFB) *in vitro*, load the cells with magnetic nanoparticles and inject these into infarcted mouse hearts using magnetic steering. This yields strongly increased numbers (∼4 fold compared to other cell types) of engrafted mFB. The injected mFB and endogenous myofibroblast (endoFB) population remain separate in the scar, but grafted mFB enhance the proliferation rate of endoFB by ∼4 fold. We also tested the functional impact of this approach by grafting lentiviral (LV)-transduced Connexin43 (Cx43) overexpressing mFB into the cardiac lesion. Prominent engraftment of Cx43^+^ mFB provides strong protection against post-infarct ventricular tachycardia (VT) *in vivo*, as VT incidence is reduced by ∼50 % at two and eight weeks after cell injection. Thus, *ex vivo* gene and subsequent *in vivo* cell therapy combined with magnetic steering of cardiac mFB enable efficient functional targeting of the cardiac scar.

## 2. Introduction

Cardiovascular disease and in particular myocardial infarctions (MI) are among the most frequent causes of death worldwide. Common and potentially life-threatening complications of the acute and chronic phase of severe ischemic heart disease are VT (1, 2). Unfortunately, besides devices, such as implantable cardioverter defibrillators (ICD), there are currently no causal therapies available to treat and improve the long-term outcome of VT. One reason is that the myocardial scar has so far proven particularly challenging for gene therapy-based targeting approaches. In fact, the most widely used viral vectors in rodents and also humans, such as adeno-associated (AAV) and lentivirus (LV), have a low efficiency to transduce cardiac mFB in the scar *in vivo* (3). Likewise, also cell therapy is challenging, as numbers of grafted embryonic cardiomyocytes (eCM) strongly decline over time upon injections into the cardiac lesion (4–6). We have shown that this is due to the retrograde flux of eCM out of the injection channel and to cardiomyocyte (CM) apoptosis, most likely because of the adverse local conditions. We have successfully improved engraftment rates of eCM by loading the cells with magnetic nanoparticles (MNP) and magnet-assisted injection into the lesion (6, 7). Given the poor results with viral targeting of the cardiac scar and our success with cardiac cell grafting, we explored the utility of combining *ex vivo* gene therapy, MNP loading and magnet-assisted cell injection into cardiac lesions as a novel cardiac targeting and treatment strategy. Cardiac mFB are a large cell population in the mammalian heart, they can be harvested and expanded *in vitro* and are reported to be more tolerant towards the adverse local conditions (8–10). Moreover, for translational purposes human cardiac mFB can be obtained from donors or generated from human induced pluripotent stem cells (hiPSCs) (11–13). We therefore explored the utility of the combined *ex vivo* gene and *in vivo* cell therapy approach with magnetic steering, and found strongly enhanced engraftment rates of mFB. We have identified PMAO-MNP as the best suited particles. We have also probed the functional impact of this strategy by overexpressing the gap junction protein Cx43 in mFB *in vitro* and injecting the cells upon MNP loading into the infarcted mouse heart resulting in a strong anti-VT effect *in vivo* at 2-and 8 weeks post-surgery. Thus, *ex vivo* gene therapy of mFB in combination with magnet-guided grafting is a promising strategy for efficient targeting and modulating the properties of the cardiac scar.

## 3. Materials and Methods

### Generation and formulation of the PMAO-MNP

The reagents used for MNP synthesis were purchased from Sigma-Aldrich (Taufkirchen, Germany) and Merck (Darmstadt, Germany). MNP synthesis and water transfer was done, as previously described (14, 15). Briefly, oleic acid-coated 12 nm iron oxide MNP were obtained by thermal decomposition of Fe(III) acetylacetonate and coated with an amphiphilic polymer (poly (maleic anhydride-alt-1-octadecene), PMAO, MW 30000–50000 Da) having 1 % of the monomers modified with 5-carboxytetramethylrhodamine (TAMRA, AnaSpec; PMAO-MNP). Further functionalization was performed by incubating 1 mg of Fe with 66 µmol of 1-ethyl-3-(3-dimethylaminopropyl) carbodiimide (EDC) and 80 µmol of EDC and 15.4 µmol of 4-aminophenyl β-D-glucopyranoside; Glc-MNP) in 250 μL of SSB buffer pH 9 (50 mM of boric acid and 50 mM of sodium borate). After three hours of reaction (stirring at room temperature and protected from light), the excess of reagents was eliminated by performing several washing steps with Milli-Q water using 4 mL cellulose membrane centrifugal filters (Amicon, MilliPore, 100 kDa, Merck). All MNP suspensions were sterilised by filtration using 0.22 μm MilliPore® filters before addition to cell cultures. Hydrophobic 12 nm iron oxide MNP were obtained in two steps by thermal decomposition of iron acetylacetonate, as previously described. The MNP were then transferred to aqueous phase using the amphiphilic polymer poly (maleic anhydride-alt-1-octadecene) (PMAO). To enable *in vitro* and *in vivo* tracking of the MNP, the polymer was modified prior to the water transfer step with 5-carboxytetramethylrhodamine (TAMRA).

### Lentivirus (LV) production and titration

The preparation, purification and titration of self-inactivating LV-vectors (rrl-CMV-IRES-eGFP or rrl-CMV-Cx43-IRES-eGFP) were performed, as described in detail earlier (3, 16).

### LDH and MTT Toxicity Assays

NIH-3T3 fibroblasts (3T3-FB, 1.0×10^4^ cells) were cultured in 10 % FCS/DMEM (Gibco, Life Technologies, Life Technologies, Carlsbad, CA, US). 3T3-FB were incubated with MNP (E-SO-2-MNP, SoMag5-MNP (6), Glc-or PMAO-MNP were added to cell culture medium) in concentrations ranging from 5 pg - 100 pg Fe/cell for 60 minutes with or without applying a magnetic field via a disk magnet positioned at 5 mm underneath the cell culture dish. The toxicity of MNP was determined 24 h upon MNP treatment using the LDH-toxicity Assay (Promega Corp, Fitchburgh, WI, US) according to the manufacturer’s instructions. Results are given in toxicity levels [%] per pg Fe/cell. Proliferation rate of NIH-3T3 fibroblasts was determined 24 h after MNP treatment via MultiTox-Fluor Multiplex Cytotoxicity Assay (Promega). As controls, fibroblasts were either untreated or treated with 2 % DMSO.

### Magnetic Particle Spectrometry (MPS)

For MPS 4.0×10^4^ NIH-3T3 fibroblasts were seeded onto 24-well plates and cultured in 10 % FCS/ DMEM (Gibco). Cells were treated for 60 min (at 37 °C and 50 rpm) with MNP (added to cell culture medium) in different concentrations (15, 25, 50 pg Fe/cell) with or without application of a magnetic field. Afterwards, cells were detached via a 5 min exposure to Trypsin (Gibco), cell numbers were determined and cells were transferred into a particulate-free reaction tube containing 2.5 % agarose gel. MPS analysis was done in cooperation with PTB Berlin, as reported before (6, 17, 18).

### Analysis of retained cell number post magnet application *in vitro*

In order to determine the retention of MNP-treated cells, NIH-3T3 fibroblasts were seeded with a density of 2.5×10^5^ cells onto a 6-well plate. After 24 h cells were treated with MNP by using either 25 or 100 pg Fe/cell iron concentration. After additional 24 hours, cells were detached with Trypsin (5 min at 37 °C) and the total cell number counted. Thereafter, cells were transferred onto a magnet rack (customised, laterally mounted magnet) and numbers of retained cells were determined, as described before (17, 19).

### Lentiviral (LV) transduction of fibroblasts with or without MNP-complexation

For PMAO-MNP two different transduction protocols were tested using NIH-3T3 fibroblasts: A two-step protocol was used, where 7.5×10^3^ cells (seeded on plates) were first incubated overnight with the LV-constructs (MOI=5) and three days later, also overnight, loaded with MNP (25 pg Fe/cell), with or without (control) magnetic field application. Alternatively, the LV were complexed with the MNP in HBSS^++^ at RT for 20 min (LV MOI=5, PMAO-MNP 25 pg Fe/cell). Then, adding the LV/MNP complexes for 30 minutes under magnetic field application at 37°C, the fibroblasts were loaded with MNP and simultaneously transduced with LV (19). Three days after transduction, the cells were fixed and analysed by immunohistochemistry.

### Isolation, culturing, LV transduction and MNP loading of murine embryonic cardiac (mFB)

Hearts of mouse embryos (E13.5, CD1 wild type) were harvested and enzymatically digested for 45 min at 37 °C with collagenase type II (520 U/ml, Worthington, NJ US) (20). The isolated cells were plated (0.1 % gelatine-coated T75 flasks, approx. 1.0×10^7^ cells) and cultured in 10 % FCS-containing DMEM (Gibco). Medium was changed every 3 days. At confluence, cells were detached using Accutase (approx. 5 min at RT, Merck) and further passaged (one passage at d3, 1:2 splitting). To obtain sufficient cell numbers for a series of transplantation experiments, cultivation time of the cells was at least 12 days, at this time the cell population was cardiomyocyte-free.

For transplantation experiments, mFB were transduced overnight (at d8) with the LV-constructs (MOI=5), as described for NIH-3T3 fibroblasts and then seeded onto 6-well plates coated with 0.1 % gelatine (1.0×10^6^ cells per well). 24 h prior to intramyocardial injections, PMAO-MNP were added to the cell culture medium overnight at 37 °C at a concentration of 25 pg Fe/cell (at d11). Prior to surgery, MNP-loaded mFB were detached with Accutase (10 min, RT) and washed at least 3 times with PBS, with 5 min of centrifugation at 1000 rpm each time. Finally, for preparation of the injection solution, mFB were dissolved in 10 % DMEM at a concentration of 0.4×10^5^ cells/µl. Then, 5 µl of this solution containing a total amount of 2.0×10^5^ mFB were injected intramyocardially into each animal (at d12).

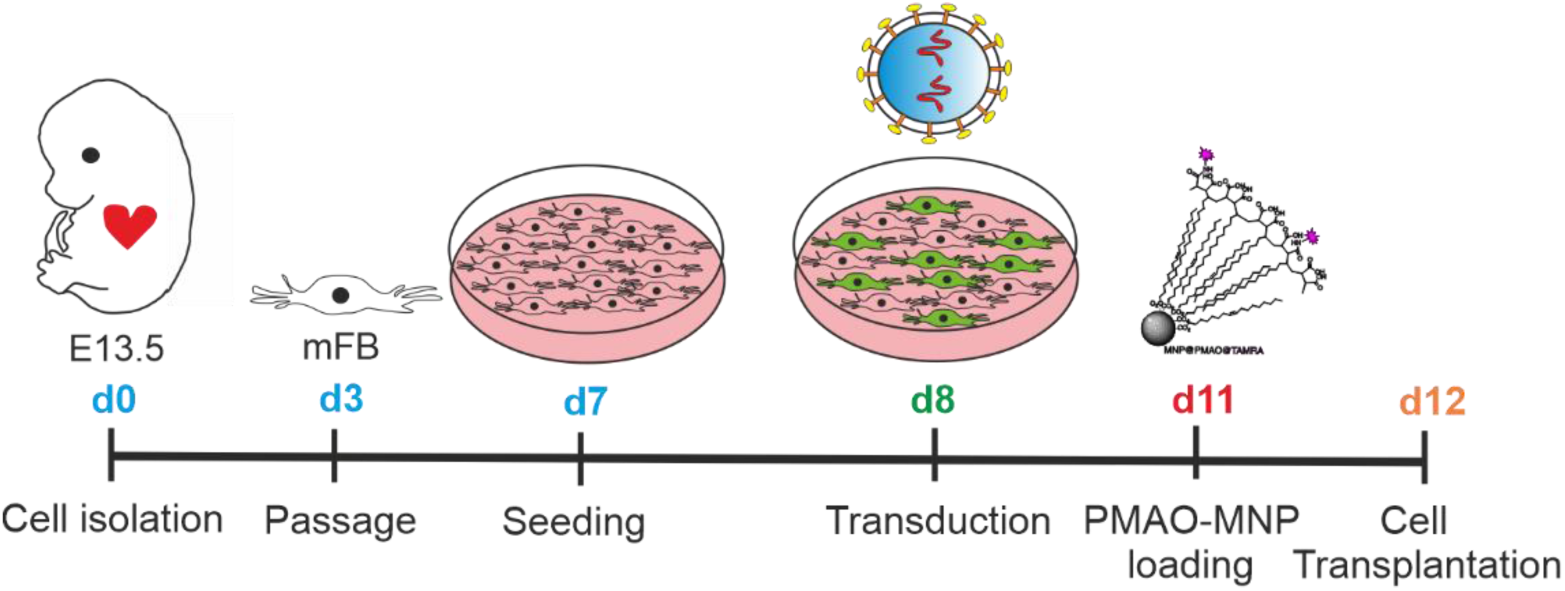
Timeline of mFB isolation, culturing, lentiviral transduction and PMAO-MNP loading prior to grafting.

### Evaluation of transduction efficiency by Flow cytometry

Flow cytometric analysis of LV-transduced mFB was performed on day 3 post transduction. Briefly, cells were detached with Accutase, washed with PBS and resuspended at a concentration of 5×10^5^ cells in 450 µl FCS-free DMEM. Shortly before analysis, 50 µl of Hoechst dye (Sigma-Aldrich) was added and samples were analysed using a FACSAria Fusion (BD Bioscience, Franklin Lakes, NJ, USA). Gating and relative quantification were performed using the FlowJo V10 software (BD Bioscience, Franklin Lakes, NJ, USA).

### Isolation of murine neonatal cardiac transgenic myofibroblasts (exoFB) for transplantation

Transgenic exoFB for grafting experiments were obtained from culturing FB of crossbred of the double-transgenic two-colour fluorescent Cre-reporter line: mTmG (Gt(ROSA)^26Sortm4(ACTB-tdTomato,-EGFP)Luo^) (21) x Tcf21^MCM^ mice at P3 (mixed background of CD1 and Bl6). For Tcf21-eGFP induction (fibroblast labelling), neonates were treated daily with Tamoxifen (intragastric injection, 0.03 mg/g body weight) from P0 to P2 (3 doses in total). Neonatal hearts were harvested at P3 and mFB isolated using collagenase type II. exoFB were cultivated in 10 % FCS containing DMEM for 3 days and then passaged and seeded onto 0.1 % gelatine coated 6-well plates. On day 4 of cultivation, exoFB were treated overnight with MNP in cell culture medium (PMAO-MNP, 25 pg Fe/cell). On day 5, exoFB were detached and prepared for cell injections (as described above). The idea was that all eGFP^+^ cells are (m)FB, however, it turned out also as reported before (22), recombination was in the range of only 60-80 %. Furthermore, our histological analysis of the tomato^+^ cell fraction revealed that < 90 % of tomato^+^ cells were positive for mFB markers (Suppl. Fig 4a). Therefore, in later experiments the pubs were not treated with tamoxifen, as all cardiac cells are tomato^+^. The engraftment rates of cells were quantified by either counting eGFP^+^/tomato^+^ cells (after tamoxifen treatment) or tomato^+^ cells (no tamoxifen treatment). For simplicity reasons, mFB are depicted exclusively in red fluorescence using pseudo-colour.

### Grafting of mFB or exoFB into infarcted mouse hearts

For grafting, first a cryolesion (CI) was generated (see Fig below) in adult (10-12 weeks) CD1 wildtype female mice, as reported before (3, 6, 7, 20): Briefly, a left lateral thoracotomy was performed under inhalative anesthesia (1-2 Vol. % Isoflurane (IsoFlo®, Zoetis GmbH, Munich, Germany) and 0.8 L/min oxygen) and a copper probe (precooled in liquid nitrogen, 3.5 mm diameter) was adhered 3 times for 15 s each onto the left anterolateral ventricular wall. Then, 2.0×10^5^ lentiviral transduced and MNP loaded mFB or exoFB were injected into the center of the CI using a 10 µl Hamilton syringe equipped with a 29 G needle. During and for 10 minutes after the injection a bar magnet was positioned with a micromanipulator at 5 mm distance to the surface of the heart (see also (6)). Then, the chest was closed, de-aired and the mice allowed to wake up. Mice received peri-and postoperative analgesia (0.1 mg/kg Burprenorphin, Burprenovet, Bayer AG, Leverkusen, Germany) s.c. pre-operatively and 3 times daily up to 3 days post-surgery and in addition, overnight diluted in drinking water up to the third postoperative day. Mice also received Prednisolon (1 mg/kg diluted in 5 % glucose i.p., Merck) up to 10 days post-surgery to reduce the potential immune response to the LV-vectors.

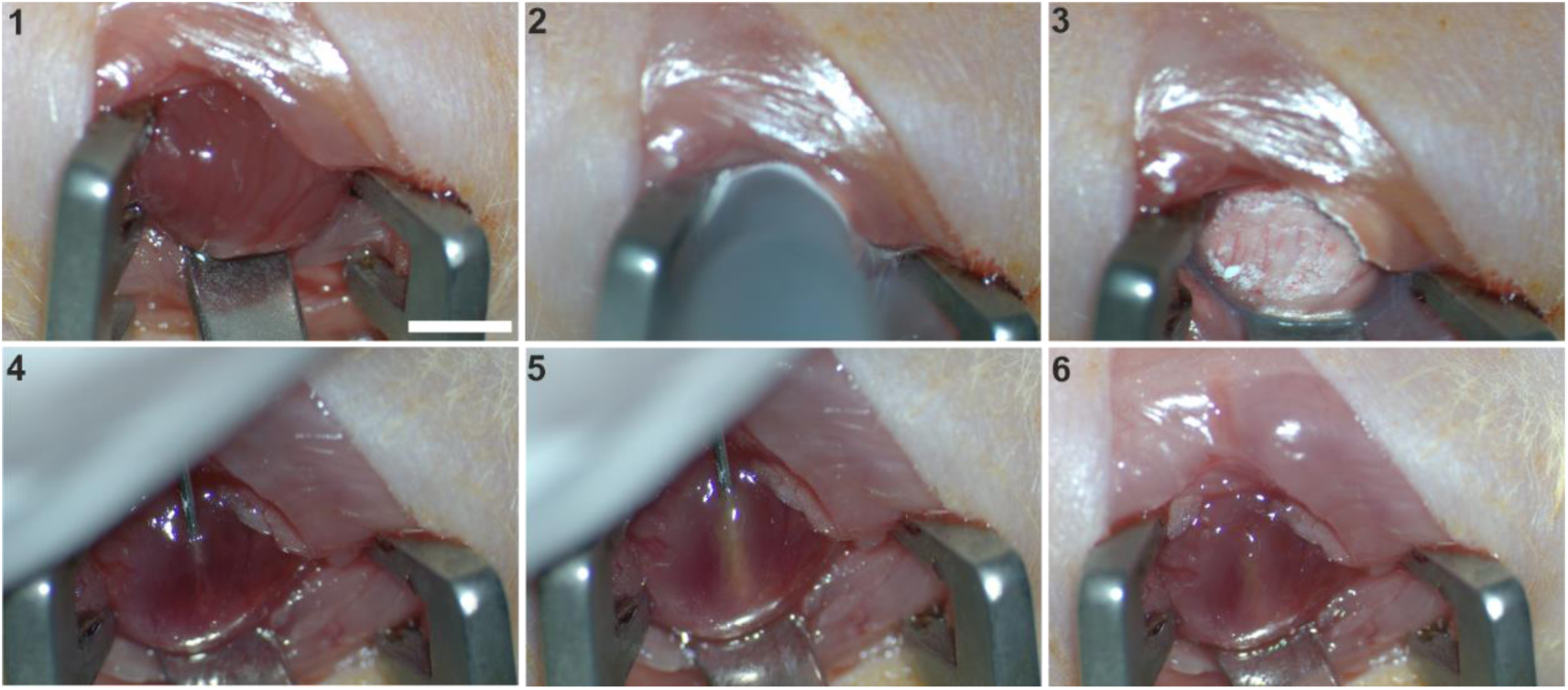
Scheme showing cryoinjury (CI) and magnet-assisted injection of mFB. 1= left sided thoracotomy, 2= cryoinjury with precooled copper stamp with a diameter of 3.5 mm, 3= freshly and frozen infarction area, 4 and 5= injection of mFB with a 29 G needle into the central part of the lesion, 6= removal of the injection needle. bar= 5 mm.

### Functional analysis with echocardiography and electrophysiology *in vivo*

Transthoracic echocardiography was performed at 2-and 8 weeks post-surgery, one day before *in vivo* electrophysiology testing. Left ventricular function was measured under inhalative anesthesia using a Phillips CX50 ultrasound system (ATL-Phillips, Oceanside, CA, USA) equipped with a 15-MHz transducer in short-axis-M-mode recordings at the level of the papillary muscles.

*In vivo* electrophysiology testing was performed at 2-and 8 weeks post-surgery under inhalative anesthesia (day 14 and 56), as reported earlier (3, 7). Briefly, the tip of a 2 French octapolar mouse-electrophysiological catheter (CIBER Mouse Electrophysiology Catheter, NuMed Inc., Hopkinton, NY, US) was inserted into the apex of the right ventricle via right jugular vein while a surface 6-lead ECG was recorded (PowerLab 16/30, LabChart 7, ADInstruments, Pty LTD, Australia). Additionally, bipolar intracardiac electrograms were recorded on atrial, his-bundle and ventricular levels. Electrophysiological vulnerability was tested in extra-and burst stimulus testing. Rectangular stimulation pulses were applied via the apical pair of electrodes by use of a multiprogrammable stimulator (Model 2100, A-M Systems, USA; Stimulus 3.4 software, Institute for Physiology I, University of Bonn, Germany).

Extrastimulus stimulation was performed with up to 3 additional beats (with stepwise S1S2 or S2S3 reduction by 10 ms, starting 10 ms below S1S1) at S1S1 cycle lengths of 120 ms, 100 ms, and 80 ms. During burst stimulation, S1S1 stimulations with cycle lengths starting at 50 ms and gradually decreasing to 10 ms were each performed for 1 second and repeated three times. Between stimulation events, the hearts were allowed to recover for at least 4 s. According to the clinical definition and previous publications, a VT was characterised by at least four consecutive ventricular extra beats with atrioventricular dissociation (23).

### Immunofluorescence stainings

For *in vitro* immunostainings, cells were seeded on glass slides (1.0×10^4^ cells/well) in 24-well plates coated with 0.1 % gelatine and fixed with 4 % paraformaldehyde (PFA, Sigma-Aldrich) for 20 min at RT. The different antibodies were added at concentrations indicated below and diluted in a volume of 80 µl per glass slide. Macroscopic native and fluorescence images of harvested hearts were acquired with a Zeiss Axio Zoom V16 macroscope (Carl Zeiss Microscopy GmbH, Jena, Germany). Hearts were fixed overnight at 4 °C with 4 % PFA and incubated for further 24 h with 20 % sucrose solution, before they were embedded in Tissue-Tek® medium (Sakura Finetek, Europe B.V., Umkirch, Germany) and frozen on dry ice. From the frozen hearts, 7 µm thick sections were prepared using a cryotome (Leica Biosystems, Wetzlar, Germany). To detect the transplanted mFB and to analyse Cx43 expression in the scar area, the sections were permeabilised with 0.2 % Triton-X-PBS (Sigma-Aldrich) and simultaneously blocked with 5 % donkey serum (DS) (Jackson ImmnuoResearch, Ely, UK). Then, antibodies (Cx43: 1:800 custom-produced rabbit polyclonal; αSMA: 1:800 mouse monoclonal; Vimentin: 1:1000 chicken polyclonal; CD45: 1:800 rat monoclonal (all Merck); CD31: 1:400 rat monoclonal (Becton Dickinson GmbH, Heidelberg, Germany); Ki67: 1:200 rabbit monoclonal; cTnT: 1:200 mouse monoclonal (both Thermo fisher Scientific GmbH, Dreieich, Germany); PDGFRα: 1:200 goat polyclonal, RD Systems;, Thermo fisher; clCaspase3: 1:200 rabbit, Cell Signalling Technology, Cambridge, UK; αActinin: 1:400 mouse monoclonal (Sigma-Aldrich)) were diluted in 5 % DS/PBS and applied for 2 h at RT to the tissue sections, followed by 3 washing steps using PBS. Secondary antibodies (all 1:400, Jackson ImmunoResearch) were diluted in Dapi (Thermo fisher) and applied for 1 h at RT. After washing, slides were covered aqueous (Aqua Polymount, Thermo Fisher) and imaged using confocal microscopy (Eclipse Ti, Nikon Europe B.V., Amstelveen, Netherlands) or fluorescence microscope (BZ-X810, Keyence, Neu-Isenburg, Germany).

### Analysis of infarction size

Infarct surface was determined using macroscopic images of the infarcted hearts acquired with an Axio Zoom V16 (Zeiss, 8x magnification). The infarct borders were clearly visible due to the CI, and the surface area (given in mm^2^) was measured using ImageJ software (NIH, Bethesda, MD, US). For morphometric analysis of infarct volumes, hearts were cryopreserved and sectioned as described above. Over the entire heart, every 300 µm sections were analysed and the measured infarct area extrapolated to the total volume of the myocardium. To visualise collagen-rich tissue and thus clearly mark the scar areas, the heart sections were stained with Sirius red (Sigma-Aldrich) and Fast Green (Fisher-Scientific). Overview images (32x magnification) were taken with a macroscope (Zeiss, Axio Zoom V16) and analysed with ImageJ. Infarct volume was expressed as % of total left ventricular volume.

### Quantification of engrafted fibroblasts

For quantification of engrafted mFB, hearts were cryopreserved and serially cut (7 µm thickness, as described above). To better visualise the eGFP fluorescence of transduced embryonic mFB, slides were incubated with anti-eGFP (1:200 goat polyclonal, Santa Cruz Biotechnology, Delaware, US) after antigen retrieval. In addition, slides were incubated with anti-Vimentin (1:1000 chicken polyclonal, Merck) as fibroblast marker (see staining protocol above) and nuclei with Dapi (Thermo fisher). Images were recorded using an Axio Observer microscope equipped with epifluorescence (Zeiss). The number of engrafted eGFP^+^ or tomato^+^ and Vimentin^+^ mFB was determined in 3 hearts per experimental group. For this purpose, the labelled mFB were counted in a total of 8 slices with a distance of 280 µm per infarct area including the border zone and extrapolated to the total volume of the myocardial lesion.

### RNA isolation, quantification and RNAseq analysis of samples

RNA from mFB (see above) was isolated (according to manufacturer instruction) by using the RNeasy© Mini Kit (Qiagen, Hilden, Germany). After RNA quantification by absorption measurement, RNA quality was determined with a Bioanalyzer (Agilent Technologies, Waldbronn, Germany) using the RNA 6000 Nano Kit (Agilent). For RNA-Seq and qPCR analysis, RNA samples with a RIN (RNA integrity number) >7 were used, the RIN’s of analysed samples ranged between 9-10. A RNA library was generated according to manufacturer instruction by use of the Trio RNA Seq library Kit (NuGEN Technologies Inc, San Carlos, CA, US). RNA bulk sequencing was done (EMBL, Heidelberg, Germany) by paired-end sequencing including 10-15 Mio reads.

### Gene expression analysis using TaqMan™ qPCR

The gene expression in transduced mFB was investigated by qPCR using TaqMan assays (Thermo Fisher). For this, 250 ng RNA was transcribed into cDNA using the SuperScriptVILO Kit (Thermo Fisher). Then, 2 µl cDNA was amplified with TaqMan™ qPCR (38 cycles, 3 technical replicates of 3 biological replicates) using master mix (Thermo Fisher) and the following TaqMan assays: eGFP (customized), Cx43 (customized), Gata4 (#Mm01310448_m1), Tnnt2 (#Mm00441920_m1), Atp2a2 (#Mm01201431_m1) and Caspase1 (#Mm00438023_m1) (all assays: Thermo Fisher, FAM). ΔCT values were calculated with mean sample CT normalized to mean CT values of the endogenous housekeeping gene HPRT (Thermo Fisher, #Mm03024075_m1, VIC) detected via Multiplex qPCR.

### Western blotting

Protein isolation of *in vitro* samples was performed using RIPA buffer (50 mM Tris-HCL pH 7.5, 1 % NP-40, 0.25 % Na-Desoxycholate, 150 mM NaCl, 1 mM EDTA pH 8.0, 1mM PMSF, Pierce™ Protease& Phosphatase Inhibitor Mini Tablets) and final protein concentration was determined with BCA protein Assay (Biorad Laboratories, Feldkirchen, Germany). Heart tissue samples of the infarct areas were excised (2 weeks or 8 weeks post-surgery) using a macroscope equipped with fluorescence (Axio Zoom V16, Zeiss). For protein isolation, heart tissue was homogenized (with micro pellet pestle (Wilmad LabGlass (Vineland, NJ, USA)), at 4°C) using RIPA buffer (see above) and protein concentration was determined as described above. For gel electrophoresis, SDS mini gels (8 % or 12 % separating gel: 8 % or 12 % Acrylamide, 7.5 mM Tris-HCL (pH 8.8), 0.1 % SDS, 0.05 % APS, 0.01 % TEMED; 4 % or 8 % stacking gel: 4 % or 8 % Acrylamide, 1.25 mM Tris-HCL (pH 6.8), 0.1 % SDS, 0.05 % APS, 0.01 % TEMED) were prepared, proteins diluted in 2x or 4x Lämmli buffer (Biorad) (for 2x Lämmli: 1:2 dilution, for 4x Lämmli: 1:4 diution) to 20 µg and denaturated for 10 min at 100 °C. Proteins were separated by electrophoresis (ProSieve EX Running buffer 10x, diluted in H_2_O to 1x, (Lonza, Basel, CH); Precision Plus Protein WesternC Standards, Biorad). Then, separated proteins were blotted onto PVDF-membranes (Methanol activated, Low fluorescence, 0.2 µm (Biozym Scientific GmbH, Hessisch Oldendorf, Germany)) by tankblot for one 1.15 h and 100 V (ProSieve EX Transfer buffer 10x, diluted in H_2_O to 1x, Lonza). Membranes were blocked with 5 % milk powder (MP, Skim milk powder, (VWR, Radnor, PA, US)) in 1x TBST buffer (20 mM Tris, 150 mM NaCL, 0.1 % Tween-20 in H_2_0) for minimum 1 h. Antibodies, diluted in 5 % MP/TBST or 5 % BSA/TBST, (Cx43: 1:3000 custom-produced rabbit polyclonal; eGFP: 1:1000 mouse monoclonal (Clontech Laboratories Inc, Mountain View, CA, US); αSMA: 1:1000 mouse monoclonal, Merck; Vimentin: 1:1000 rabbit monoclonal (Cell signalling); Periostin: 1:1000 rabbit polyclonal (Novus Biologicals, Centennial, CO, US); TGFb1: 1:500 rabbit polyclonal (Abcam, Cambridge, UK) were incubated overnight at 4 °C. As housekeeper, Horseradish peroxidase (HRP) conjugated-GAPDH (1:5000, Sigma Aldrich) or ß-Actin-HRP (1:10000, Sigma-Aldrich) was incubated for 1 hour at room temperature and in 5 % MP/TBST. Followed by 3 washing steps with 1xTBST, secondary antibodies (donkey-anti-rabbit Alexa Fluor 488: 1:3000, Affinipure, Jackson Immuno Research; donkey-anti-mouse IGg2a Cy5; donkey) and Precision Protein StrepTactin-HRP Conjugate (1:3000, Biorad), diluted in 5 % MP/TBST, were incubated for 1 hour at room temperature. Signals were developed with Pierce ECL Western Blotting Substrate (Thermo Fisher) and detected with ChemiDoc MP Imaging System (Biorad). Quantification of target protein expression was performed with Image Lab Software (Biorad) and normalised to GAPDH or ß-Actin expression.

### Statistics

Electrophysiological data were compared using a Fisher’s exact test. Statistical evaluation of data with more than two groups was performed using one-way Anova with posthoc Turkey test. Data with two groups was analysed using Student’s t-test. A p value ≤ 0.05 was considered statistically significant, error bars represent SEMs.

### Study approval

All mouse experiments were performed in accordance with the ARRIVE guidelines, the guidelines of the German law of protection of animal life and approved by the local government authorities (Landesamt für Natur und Verbraucherschutz Nordrhein-Westfalen, NRW, Germany). Animal protocol numbers: 81.02.04.2018.A120 and 81-02.04.2020.A093.

## 4. Results

### Identification and characterisation of best suited MNP

In our earlier study we showed strongly improved engraftment rates of CM upon combining SoMag5-MNP with magnetic steering (6). However, these MNP formed aggregates in the cytosol and displayed cellular toxicity (Fig 1e,f). We, therefore, tested different types of MNP focusing on low toxicity and high magnetic potential for efficient loading and transduction of (m)FB. We tested 4 MNP types with different chemical and biophysical properties (E-SO-2-MNP, Glc-MNP, SoMag5-MNP, PMAO-MNP) in NIH-3T3 fibroblasts (3T3-FB). The combination of MNP and magnetic field application (24 well plate, Chemicell, Berlin, Germany) for 60 min increased cellular toxicity at concentrations > 25 pg Fe/cell compared with MNP without magnet (Fig 1a,b; left panels). PMAO and SoMag5-MNP displayed the lowest toxicity even at high Fe concentrations in presence of a magnet (100 pg Fe/cell) (Fig 1a, right panel) in the LDH assay, whereas the other 2 MNP types were more toxic (Fig 1a, right panel, 50-70 % toxicity at 100 pg Fe/cell+magnet). The MTT-assay yielded similar results (Fig 1b), as 58 %-and 54 % viable cells were found at 25 pg Fe/cell for PMAO-and SoMag5-MNP, respectively, whereas the other particles yielded poor results (see also Fig 1b). To exclude that the decreased toxicity of SoMag5-and PMAO MNP was due to lower cellular Fe uptake we measured the Fe content using magnetic particle spectrometry (MPS). This revealed that cellular uptake of PMAO-and SoMag5-MNP into 3T3-FB was significantly higher compared to the other MNP (Fig 1c). Next, the magnetic potential of the MNP-loaded cells was assessed in a metal rack (Fig 1d), PMAO-and SoMag5-MNP had the highest cell retention rates with ∼70 % at 100 pg Fe/cell and ∼30 % at 25 pg Fe/ cell, respectively. Interestingly, Prussian blue staining (Fig 1f, upper panel) revealed that SoMag5-, but not PMAO-MNP displayed intracellular complex formation, of further advantage was that the intracellular distribution pattern of Tamra fluorochrome linked PMAO-MNP could be directly studied with a fluorescence microscope (Fig 1f, lower picture). In accordance with the Prussian blue data, fluorescence pictures illustrated an equal distribution of PMAO-MNP within the cytoplasm of 3T3-FB (Fig 1f, Fig 1i). The intracellular distribution pattern of the PMAO-MNP was also studied at higher resolution using a transmission microscope. This showed that PMAO-MNP (Fig 1g) formed homogeneous, non-aggregate structures with an average size of 22 nm in FB (Fig 1h). Thus, PMAO-MNP display good uptake into FB, high retention rates, low toxicity and no cytosolic aggregate formation, therefore these particles were used for all the following experiments.

**Figure 1:**
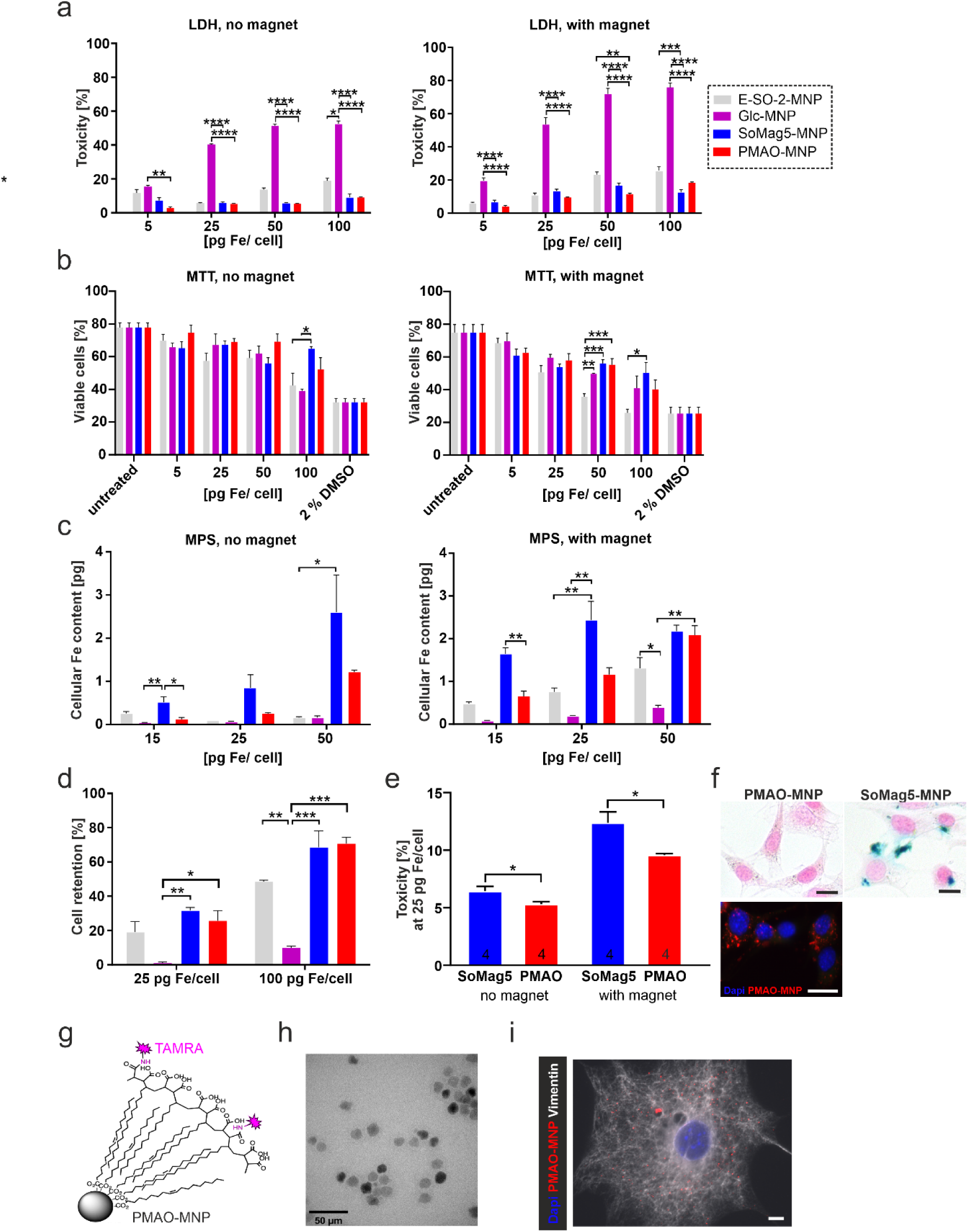
Toxicity, magnetic potential and intracellular distribution of different types of MNP in *3T3 fibroblasts*. (a) LDH-toxicity-and (b) MTT-viability-assays of MNP assessed at 24 h after MNP loading with) or without magnet application for 1h. (c) Magnetic particle spectrometry (MPS) of MNP loaded cells. (d) Retention rate of MNP loaded cells. (e) LDH-toxicity assay of 3T3 FB loaded with SoMag5-or PMAO-MNP (25 pg Fe/cell) with or without a magnetic field (f) Prussian blue iron staining of PMAO-and SoMag5-loaded 3T3-FB cells (25 pg Fe/cell: blue: iron oxide, red: nuclei, bar= 5 µm; fluorescence macroscopic image: blue= nuclei, red: PMAO-MNP, bar= 20 µm). (g) Scheme of PMAO-MNP. (h) Transmission electron microscopy image of PMAO-MNP (bar= 50 µm). (i) Microscopic image of a 3T3-FB incubated overnight with PMAO-MNP and magnet application (25 pg Fe /cell, red= PMAO-MNP, blue= nuclei, white= Vimentin, bar= 10 µm). P-values: *= < 0.05; **= < 0.01; ***= < 0.001; ****= 0.0001.

### Optimisation of transduction of mFB

We next established a protocol for lentiviral (LV) transduction and MNP loading of 3T3-FB. First, PMAO-MNP (25 pg Fe/cell) complexed with LV were added to cell culture wells and a magnetic field (30 min at 37°C) co-applied. This yielded with 21.9 % of eGFP^+^ cells a higher transduction rate compared to the 15.6 % using a two-step protocol with overnight exposure to the LV (MOI 5, 37°C) and PMAO-MNP loading at two days later (25 pg Fe/cell, 37°C, no magnet). However, the one step protocol yielded increased cytosolic MNP aggregation, whereas this was not the case using the two step protocol in 3T3-FB and mFB (Suppl. Fig 1a,b). We therefore used for the grafting experiments the two-step protocol. mFB were harvested from E13.5 mouse hearts, and propagated and enriched in cell culture. The cells were characterised using molecular and cell biological assays and we found, as reported before (24), increasing alpha smooth muscle actin (αSMA) levels during cell culture in accordance with the differentiation of cells into mFB (Suppl. Fig 1c). For transduction we used two LV, as reported before (3), one harbouring the expression cassette for Cx43/eGFP (LV-Cx43) and the control virus with only eGFP (LV-eGFP) (Fig 2a). mFB were transduced overnight at d8 after cell isolation with the LV (Fig 2a, MOI= 5) (Suppl Fig 1c) and transduction efficiency and Cx43 expression were assessed with flow cytometry yielding an efficiency of ∼15 % for both LV (Fig 2b). Next, we quantified transduction efficiency with a fluorescence microscope and found 3 days after transduction similar efficiencies for both LV with approximately 25 % (Fig 2c). The overall eGFP fluorescence was weaker upon transduction with the Cx43 LV because of the bicistronic vector construct (3). Because of the different results obtained with flow cytometry and microscopy, we counted only the bright eGFP^+^ mFB under the microscope and found that the numbers corresponded with the flow cytometry. We also quantified eGFP^+^ cells at 17 days after LV transduction, and found ∼37.6 % positive cells (Fig 2d). This could either be due that some cells require longer to accumulate enough eGFP for detection and/or a greater proliferation rate of eGFP^+^ mFB. Co-staining against Ki67 and αSMA at 17 days post-LV transduction revealed that ∼50 % of mFB were Ki67^+^ (∼48 % at d3, 52 % at d17), a slightly larger percentage of eGFP^+^ mFB (∼72 % at d3, ∼52 % at d17) were Ki67^+^ at d3 (Fig 2e). Staining against the M-phase specific marker PHH3 corroborated the prominent proliferation rate of mFB with 6.6 % and 7.5 % of native vs LV transduced mFB being positive, respectively (Fig 2f). Accordingly, cell counting yielded an increase in cell number by 4-(LV-eGFP mFB) and 3-(untreated mFB) fold, respectively, in the first 10 days post transduction (Fig 2g), proving that embryonic mFB are strongly proliferative *in vitro*. We next analysed Cx43 protein expression upon LV-Cx43 transduction of mFB using immunostaining and found that the Cx43 signal was distributed throughout the cell membrane and that it was clearly increased in eGFP^+^ mFB at 3 days after transduction (Fig 2h). This was underscored by Western Blotting, in which an approximately 2-fold increase of Cx43 protein expression in LV-Cx43 transduced compared to LV-eGFP transduced mFB was found (Fig 2i). We also investigated the impact of Cx43 overexpression in mFB by analysing the transcriptome of LV-Cx43 (n=3) and LV-eGFP (n=3) transduced mFB at 3d (Suppl. Fig 2a,b). Bioinformatic analysis revealed that 697 genes were enriched and 702 depleted in the in LV-Cx43 compared to LV-eGFP mFB (Fig 2j, left picture). Approx. 30-40 of the most downregulated genes were related to an immune response upon viral particle exposure, while the downregulation of caspase genes indicated an anti-apoptotic effect of Cx43-overexpression. When looking at specific developmental and cardiogenic genes, such as Tnnt2, Gata4, and Atp2a2 we observed an enrichment in LV-Cx43 mFB (Fig 2j, right picture). The same was true for ion channels in CM, namely Cacna1c, Kcnh2, and Scn5a as well as genes coding for transmembrane proteins (Integrinb3, Integrinb4) and angiogenesis/endothelial cell marker genes, such as Flt1, Pecam1, Gata4 and Hspg2 (Fig 2j, right picture). This data was further underscored by qPCR analysis proving that Tnnt2, Gata4 and Atp2a2 were indeed upregulated (Suppl. Fig 2c-e) indicating that Cx43 expression pushed the gene signature in the mFB towards a cardiomyogenic fate, but the cells remained positive for mFB markers such as αSMA in immunostainings (Suppl. Fig. 1d,e). Furthermore, Casp1 gene expression was significantly downregulated in LV-Cx43 mFB (Suppl. Fig 2f), confirming the results obtained with RNA-Seq. Thus, mFB show a transduction rate of approximately 40 % using LV, the cells overexpress Cx43, strongly proliferate and show changes in the gene expression pattern upon Cx43 overexpression such as adapting a signature reminiscent of CM and a downregulation of proapoptotic genes.

**Figure 2:**
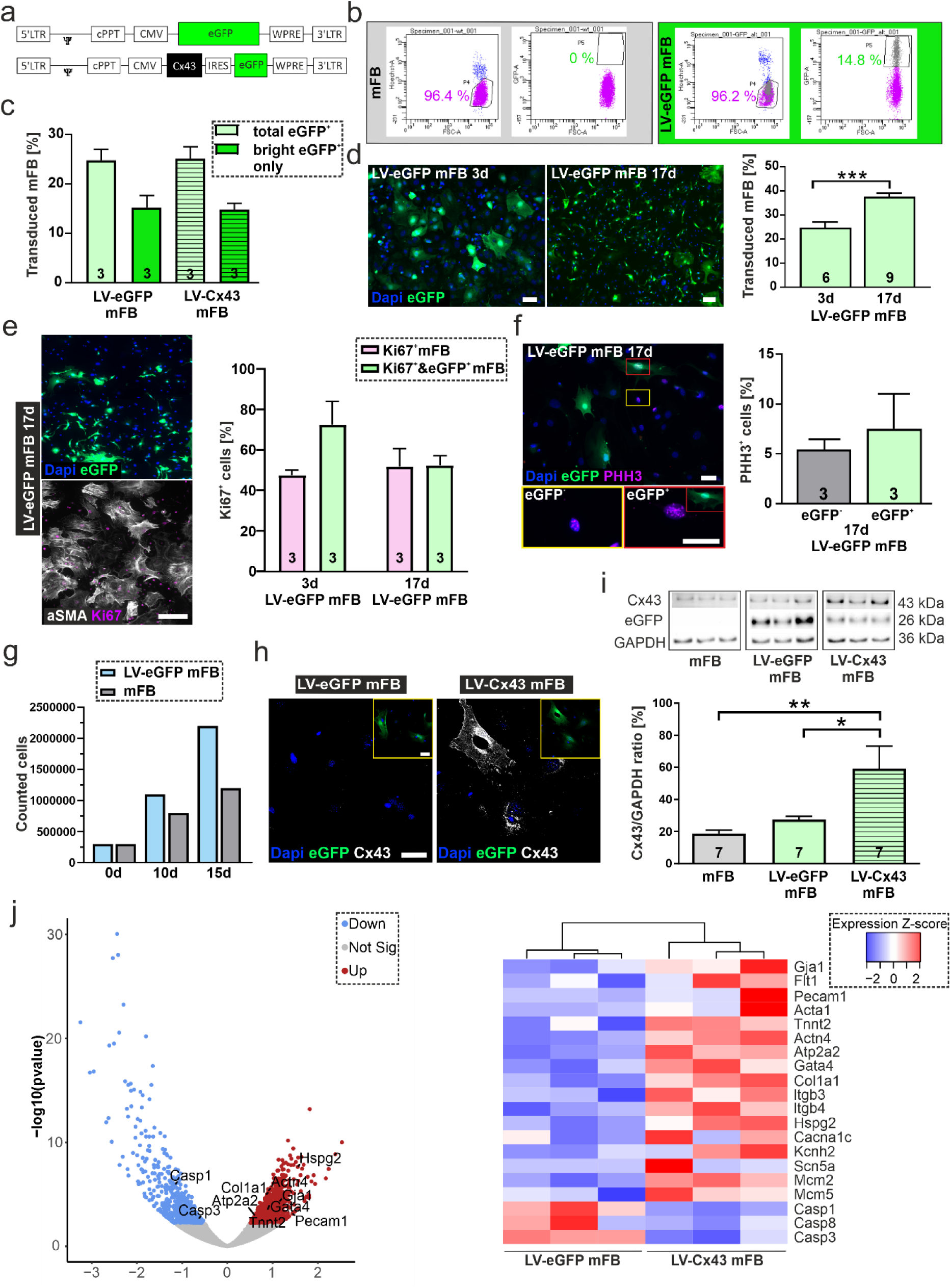
Lentivirus (LV)-based transduction of embryonic cardiac myofibroblasts (mFB) *in vitro*. (a) LV-constructs (MOI=5) used for the transduction of mFB. (b) Flowcytometric analysis (20,000 counts/sample) of LV-treated mFB (right panels) and untreated mFB (left panels). (c) Microscopic analysis of percentage total eGFP^+^ and bright eGFP^+^ mFB at 3d after overnight LV treatment. (d) Representative fluorescence pictures and percentage of eGFP^+^ mFB at d3 and 17 after LV treatment (bars= 50 µm). (e) αSMA and Ki67 co-stainings of LV-transduced mFB at 17d, quantitation of Ki67^+^/eGFP^+^ mFB at 3d and 17d post LV-eGFP transduction (bar= 200 µm). (f) PHH3 staining of LV-eGFP transduced mFB at 17d (bar= 50 µm). (g) Numbers of untreated and LV-eGFP treated mFB at different time points. (h) Cx43 overexpression in LV treated mFB at 3d (white: Cx43, blue: nuclei, green: eGFP, bar= 50 µm). (i) Cx43 protein expression at 3d in LV-Cx43 mFB, mFB and LV-eGFP mFB using Western Blotting. (j) Volcano plot and heatmap of up-and downregulated genes of LV-Cx43 vs LV-eGFP mFB at 3d obtained with RNAseq analysis (both n=3). P-values: *= < 0.05; **= < 0.01; ***= < 0.001.

### Grafting of LV-transduced mFB into the cardiac lesion

Next, we injected 2.0×10^5^ LV-transduced and PMAO-MNP loaded mFB into freshly cryoinfarcted (CI) wild-type CD1 mice (=CI, 5 µl per lesion). This injury model was used, as it is highly reproducible and best suited for the comparison of functional *in vivo* assay results between animals (3, 25). To improve engraftment rates we used magnetic steering positioning a rod magnet (1.3 T) at 5 mm distance from the surface of the heart during and for 10 minutes after injection of the MNP loaded mFB (6). Three different control groups were used, namely mice with CI, with CI and injection of LV-eGFP mFB (LV-eGFP) and magnet application and mice with CI, injection of LV-Cx43 mFB (LV-Cx43) without magnet application.

At 2 weeks after surgery, hearts were harvested and processed for histomorphological analysis. As shown in Fig 3a, in some hearts the engrafted mFB could be clearly identified with a macroscope based on the eGFP fluorescence. When grafting LV-Cx43 mFB and performing co-immunostainings for eGFP and Cx43 (Fig 3b), we found prominent cellular Cx43 expression. This was underscored by Western Blotting using excised scar tissue, in which a ∼5-6-fold increase in Cx43 protein content was detected at 2 weeks after injecting LV-Cx43 mFB *vs* controls (Fig 3c). Next, we selected LV-eGFP and LV-Cx43 hearts (both n= 3), which showed good cell engraftment under the macroscope and quantified the number of engrafted mFB after co-staining against eGFP and Vimentin (Fig 3d). We found 35,000-82,000 eGFP^+^ mFB (average = 64,000 ± 6,915 cells, n= 6), which is approximately 4-fold higher than our earlier findings injecting eCM (6). We also assessed the impact of magnetic steering by quantifying the number of Vimentin^+^/eGFP^+^ cells in LV-Cx43 hearts either with or without magnet application, the latter increased the number of engrafted mFB∼3-fold (Suppl. Fig 3); this was confirmed by Western Blotting yielding a more than 3-fold increase for both, Vimentin and Cx43 protein content. To explore, whether the proliferation of mFB could contribute to prominent engraftment rates we co-stained LV-eGFP, LV-Cx43 and control hearts against Ki67 and eGFP at 2 weeks post-surgery (Fig 3e). The number of cycling Ki67^+^ cells in the scar area was significantly higher in mFB injected hearts, as 33 % of LV-eGFP and 45 % of LV-Cx43 were Ki67^+^ (Fig 3f). Interestingly, also the number of eGFP^-^/Ki67^+^ mFB in grafted hearts was increased, but we could not determine, whether grafting had an effect on endogenous mFB (endoFB), as only 40 % of injected cells are eGFP^+^.To rule out that increased numbers of Ki67^+^/eGFP^-^ cells upon mFB injection are due to immune cell infiltration we triple-stained against CD45, Vimentin and Ki67 (Fig 3g) and found that only a very low number of CD45^+^ cells were Ki67^+^. Taken together these data demonstrate prominent engraftment and proliferation of mFB in the cardiac scar when combining MNP loading with magnetic steering.

**Figure 3:**
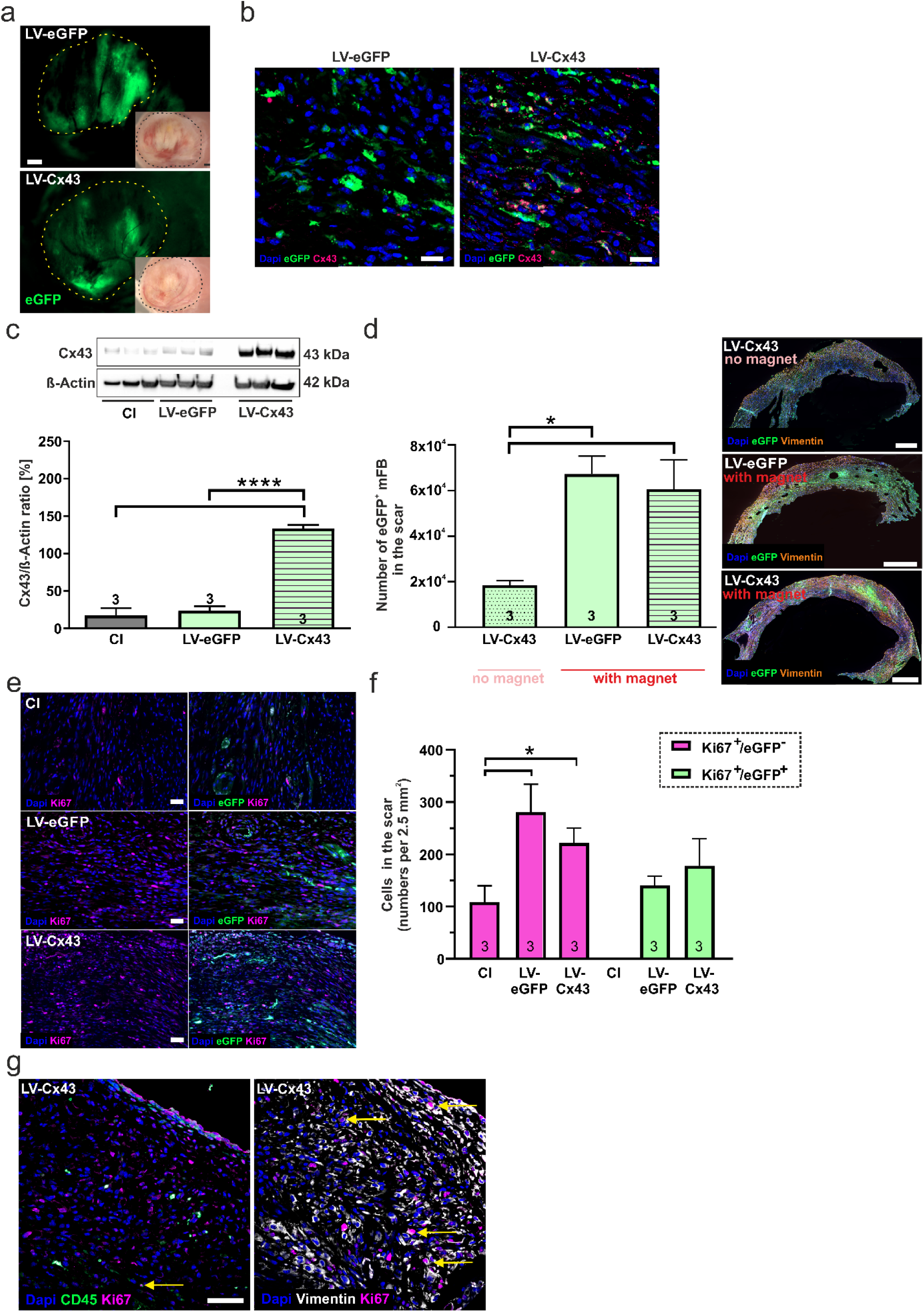
mFB engraftment and Cx43 expression in the cardiac scar of mice 2 weeks post intracardiac injection of LV-transduced mFB. (a) Macroscopic images of mFB injected CI hearts (green: native eGFP fluorescence) (dots mark the scar area, bars= 500 µm). (b) Immunostainings against Cx43 in LV-eGFP and LV-Cx43 hearts (bars= 20 µm and 5 µm). (c) Quantitation of Cx43 protein expression in cut out scars with Western Blotting. (d) Quantitation of engrafted eGFP^+^ mFB in serially and co-stained heart sections against eGFP and Vimentin, representative microscopic pictures are shown (right panels, bar= 500 µm). (e) Immunostainings against Ki67 and eGFP in scar areas of CI, LV-eGFP and LV-Cx43 hearts at 2 weeks post-surgery (bar= 50 µm). (f) Quantitation of Ki67^+^ and Ki67^+^/eGFP^+^ cells in LV-eGFP and LV-Cx43 hearts. (g) Co-immunostainings against CD45/Ki67 or Vimentin/ Ki67 (bar= 50 µm) in cardiac scars. P-values: *= < 0.05; ****= 0.0001.

### Grafting of genetically labelled neonatal mFB (exoFB) into the cardiac lesion

Because of the good engraftment rates of mFB into the cardiac lesion we reasoned that this could be the strategy to efficiently target an area which has proven so far to be quite difficult for therapeutic interventions. In order to assess more precisely the engraftment rates of injected mFB, their proliferation, distribution within the scar and interactions with endoFB, we have used double transgenic neonatal mice by crossing Tcf21^MCM^ mice with mTmG reporter mice, in which all cells are tomato^+^ and all exoFB eGFP^+^ after successful recombination. However, as recombination rate turned out to be only 60-70 %, as reported earlier (22), for most experiments P3 transgenic hearts (Fig 4a) were harvested without tamoxifen induction and exoFB enriched in cell culture. The degree of enrichment was characterised at the stage of grafting after 5 days in culture by immunostaining against αSMA showing that that >90 % were mFB (αSMA^+^, Suppl. Fig 4a); at 12 days in culture >95 % were exoFB (αSMA^+^/Vimentin^+^), and numbers of endothelial cells (CD31^+^∼4 %) and of cardiomyocytes (cTnT^+^∼0.7 %) were negligible (Fig 4b), underscoring that our cell culture protocol yielded highly enriched exoFB. The exoFB were found to proliferate, but much less than mFB, as 13 % of the cells were Ki67^+^ (Fig 4b). Cell injections were performed as described above using exoFB kept for 5 days in culture. All injected hearts were analysed without macroscopic preselection at 1-and 2 weeks (Fig 4c) post-surgery. Quantitation of exoFB yielded 53,000-92,000 (average = 72,000 ± 6,536 cells, n= 5) at 1 week and 35,000 to 74,000 cells (average = 53,000 ± 7,938 cells, n=4) at 2 weeks post-surgery (Fig 4d).

**Figure 4:**
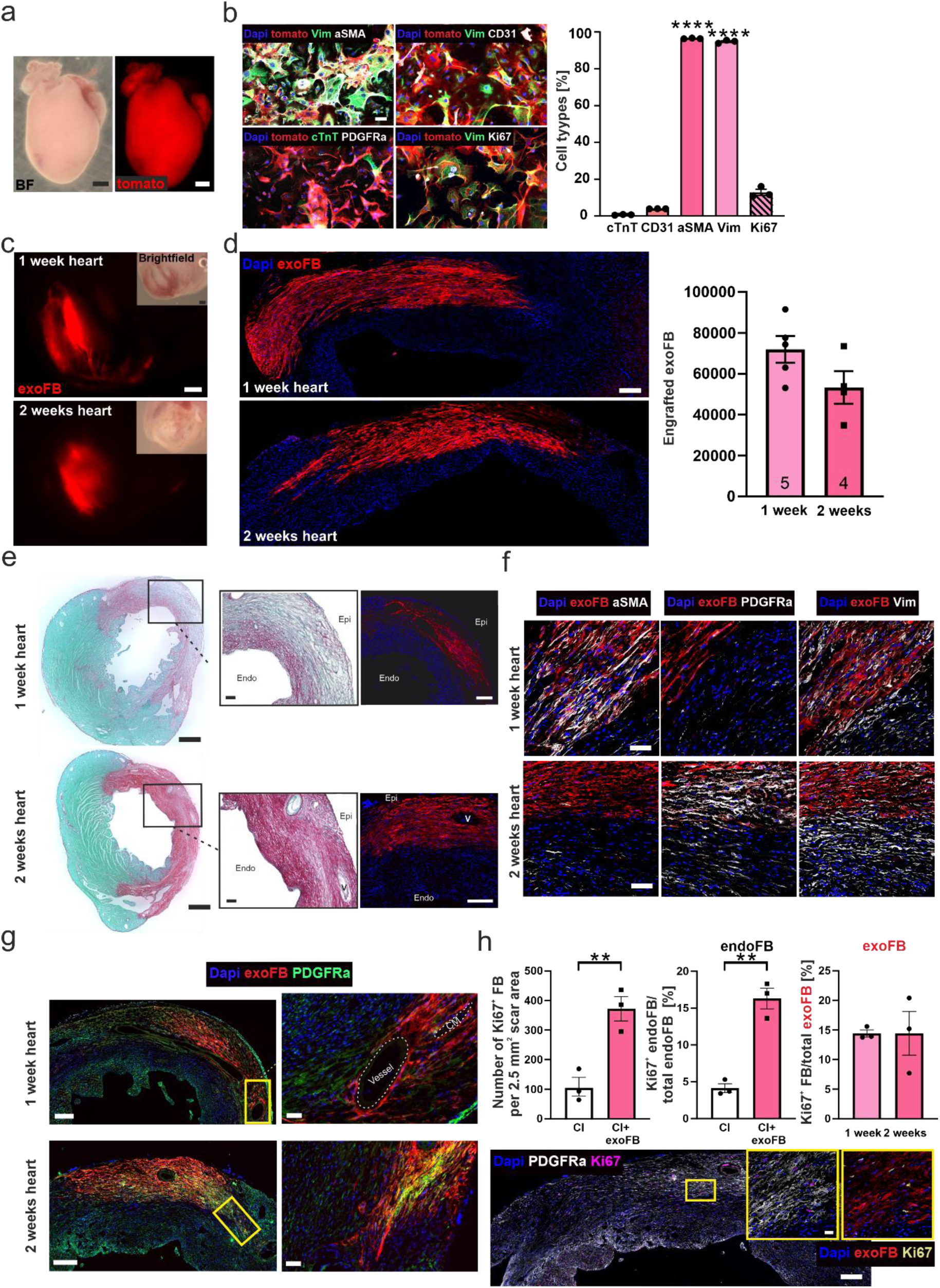
Characterisation and engraftment of neonatal cardiac mFB (exoFB) obtained from double transgenic mice. (a) Macroscopic image of tomato^+^ P3 heart (bar= 500 µm). (b) Characterisation and quantitation of exoFB and nonFB 12 days after isolation using immunostainings (blue=nuclei, red= exoFB, green= Vimentin/cTnT, white= stained marker, bar= 50 µm). (c) Macroscopic images of cardiac scar with grafted exoFB at 1-and 2-weeks post-surgery (bar= 500 µm); (d) Microscopic documentation of prominent engraftment of exoFB at 1-and 2 weeks post-surgery in the scar, quantitation of engrafted exoFB (blue=nuclei, red= exoFB, bar= 200 µm). (e) Sirius red staining of hearts 1-and 2-weeks after injection of exoFB (fluorescent images: blue=nuclei, red= exoFB, bars= 500 µm and 100 µm). (f) Immunostainings against the (m)FB markers αSMA, Vimentin and PDGFRα (blue=nuclei, red=exoFB, white=stained marker, bar= 20 µm) in 1-and 2 weeks hearts post-surgery. (g) Location and distribution pattern of tomato^+^/ PDGFRα^+^ exoFB in the scar (blue=nuclei, red= exoFB, green= PDGFRα, bars= 200 µm and 50 µm). (h) Proliferation of total FB and endoFB in CI control hearts *vs* exoFB injected hearts at 2 weeks post-surgery (left and middle graph) and proliferation of exoFB in 1-and 2 weeks post-surgery (right graph); a representative picture of an exoFB injected heart 2 weeks post-surgery is shown, boxed are is shown at higher resolution in the insets (left inset: blue=nuclei, white= PDGFRα, pink= Ki67, scale bar= 200 µm; right inset: blue=nuclei, red=exoFB, yellow= Ki67, bar= 20 µm). P-value: **= < 0.01. This figure includes a small group of transplanted exoFB obtained from Tamoxifen-induced Tcf21xmTmG P3 mice (see methods), eGFP^+^ mFB are illustrated with a red fluorescent pseudo-colour.

We next investigated the distribution pattern of the grafted exoFB in the scar, their migration and interaction with endoFB. One-week post-surgery the exoFB were preferentially found close to the epicardial layer, the cells were sticking together and located in between dying CM (Fig 4g, left upper panel). At 2 weeks post-surgery, the exoFB remained positioned close to the epicardium and sticking closer together (Fig 4g, left lower panel). Engrafted exoFB remained separate from endoFB (Fig 4g), in between these two cell populations a few CD45^+^ cells were found (Suppl. Fig 4b). The exoFB were αSMA^+^ and Vimentin^+^ proving their mFB identity, only very few tomato^+^ CD31^+^ cells were found (Suppl. Fig 4b) underscoring that almost exclusively exoFB engraft. Interestingly, the exoFB underwent after grafting further differentiation and maturation, as immunostainings revealed decreasing αSMA and increasing PDGFRα levels (Fig 4f). This was comparable to the endoFB and in line with earlier observations by other groups (26, 27). Increased collagen staining at 2 weeks post-surgery showed increasing compaction of the scar in injected and non-injected hearts (Fig 4e). We also analysed the proliferation rate of exo-and endoFB in the scar area and found significantly more Ki67^+^ FB in the scar of grafted vs CI control hearts (Fig 4h, left graph). Our analysis revealed that grafting of exoFB enhanced the proliferation rate of endoFB 4-fold, as 16.3 % of tomato^+^ endoFB were Ki67^+^ and only 4.1 % in CI controls at 2 weeks post-surgery (Fig 4h, middle graph). The proliferation rate of exoFB was with ∼10-15 % (Fig 4h, right graph) very similar to that of endoFB and to the exoFB at 12d in cell culture. Thus, the transplantation of genetically labelled exoFB showed that the cells stably integrate into the scar, and continue to proliferate and differentiate. In addition, the exoFB exert a pro-proliferative effect on endoFB.

### Functional impact of grafting Cx43 overexpressing mFB into the cardiac lesion at 2 and 8 weeks post-surgery

Given the prominent engraftment characteristics of mFB into the cardiac scar we tested whether also the functional properties of the cardiac scar could be modulated with this approach. We therefore overexpressed Cx43 in mFB *in vitro* using the LV-Cx43 vector and grafted 2×10^5^ cells into the cardiac lesion, as earlier results of our group demonstrated that Cx43 overexpression can reduce post-infarct VT incidence *in vivo* (3, 7). *In vivo* electrophysiology testing was performed using S1/S2 and burst stimulation protocols at 2 weeks post-surgery. In Figure 5a, representative traces of surface and intracardiac atrial ECG leads for control (LV-eGFP) and Cx43 (LV-Cx43) mice show that burst stimulation (Fig 5a, upper leads) induced a self-terminating VT in control hearts, whereas not VT were typically evoked in the LV-Cx43 hearts (Fig 5a, lower leads). The original traces show ventricular capture and AV dissociation during burst pacing in a representative LV-Cx43 heart, and a normo-frequent sinus rhythm after a short compensatory pause after electrical pacing (Fig 5a, lower leads). VT incidence (∼88 %) was found to be very high in CI-and LV-eGFP control mice, but significantly (CI vs. LV-Cx43, p=0.0055; LV-eGFP vs. LV-Cx43, p=0.0012) reduced in LV-Cx43 mice with ∼40 %; this incidence was very similar to the non-infarcted control group (43 %, Fig 5a). We also performed echocardiography at 13d post-surgery and surprisingly left ventricular function was improved in the LV-Cx43 group compared to controls. Cardiac pump function in M-mode recordings (Fig 5b, left graph) showed a significant (p= 0.0007 and 0.0003) improvement in fractional shortening in LV-Cx43 mice (FS= 28.9 %) compared with CI (FS= 22.2 %) and LV-eGFP mice (FS= 22.7 %), while heart rates were comparable in all three groups (data not shown). In accordance with improved left ventricular pump function we found significantly increased anterior cardiac wall thickening (ΔAWT) (p=0.0345 and 0.0315) (Fig. 5b, right graph, ΔAWT=0.31 mm) in LV-Cx43 mice compared to CI (ΔAWT=0.16 mm) and LV-eGFP (ΔAWT=0.18 mm) mice, respectively; sham operated mice showed the thickest anterior wall (0.48 mm) (Fig 5b). To exclude that the difference in LV function was due to differences in infarct size, quantitative morphometry was performed which did not evidence differences in infarct area or volume between the groups (Suppl. Fig 5a,c).

**Figure 5:**
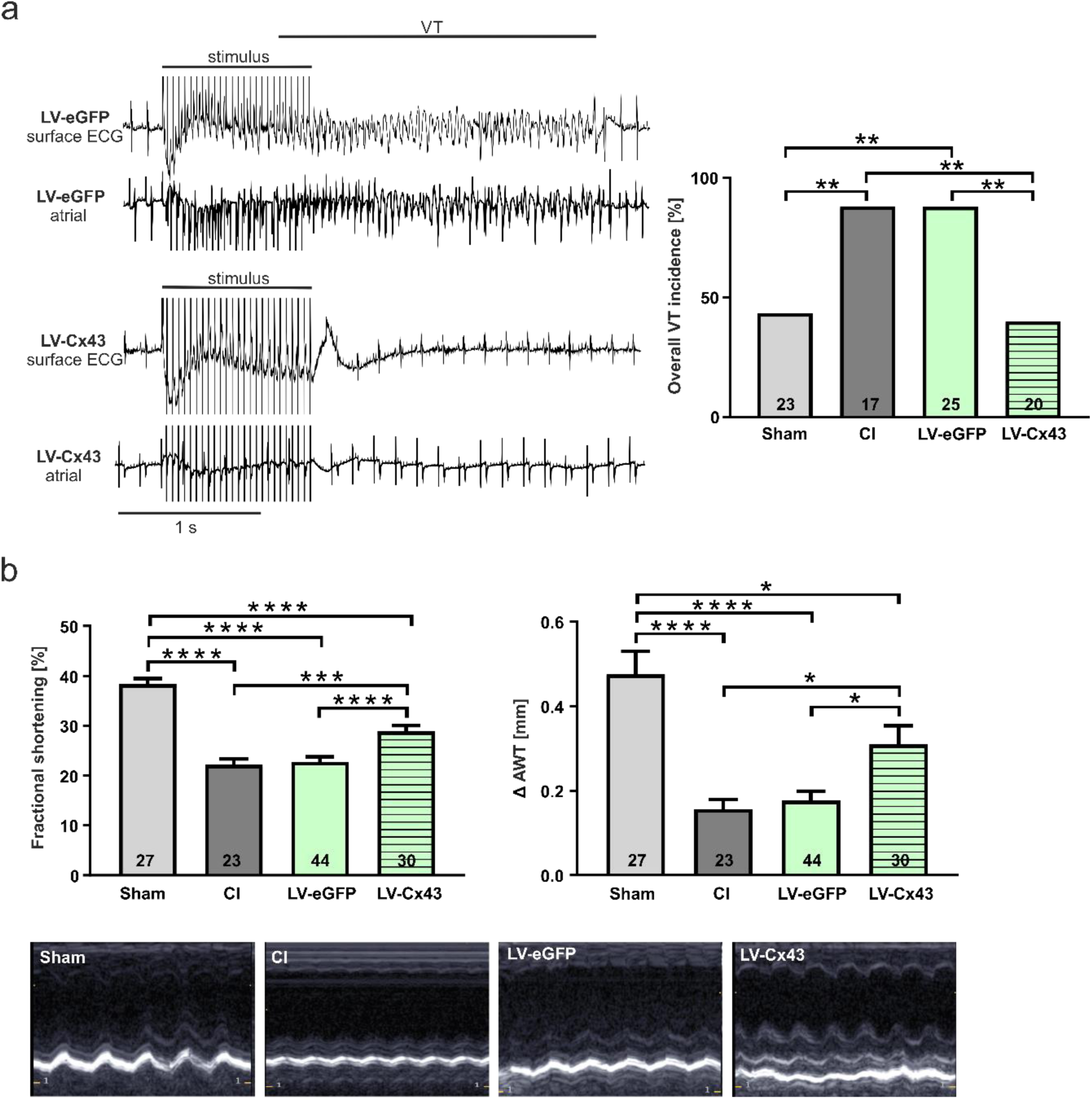
Electrophysiological testing *in vivo* and echocardiographic assessment of left ventricular function at 2 weeks post-surgery. (a) Representative surface ECG traces of a LV-eGFP (upper traces) and a LV-Cx43 (lower traces) mouse; note the self-terminating VT in the LV-eGFP mouse and capture, post-stimulation pause and sinus rhythm in the LV-Cx43 mouse upon burst stimulation at 2 weeks post-surgery. Quantitation of VT incidence in the different groups at 2 weeks post-surgery. (b) Fractional shortening (left) and anterior wall thickening (ΔAWT in mm, right). Original M-mode echocardiographic recordings from the different groups of mice. P-values: *= < 0.05; **= < 0.01; ***= < 0.001; ****= 0.0001.

We also investigated the functional effects at 8 weeks post-surgery, as it is known grafted cell numbers in the infarcted mouse heart strongly decrease over time (5, 6). At this stage we still detected with a macroscope eGFP^+^ patches in the scar indicating successful long-term engraftment of mFB (Fig 6a). We selected LV-eGFP and LV-Cx43 hearts with proven engraftment under the macroscope and found 8,000 to 39,000 (average= 17,605 ± 4,667) eGFP^+^ mFB (Fig 6d). Anti-Cx43 immunostainings in the cardiac scar showed Cx43^+^/eGFP^+^ cells in LV-Cx43 and Cx43^-^/eGFP^+^ cells in LV-eGFP hearts (Fig 6b). Cx43 and eGFP protein content was quantified in excised scars using Western blotting showing a more than threefold increase in Cx43 content in LV-Cx43 hearts compared to controls (Fig 6c). *In vivo* electrophysiological analysis revealed a significant lower VT incidence of 38.9 % in LV-Cx43 mice, compared to 88.9 % in CI and 77.8 % in LV-eGFP control mice, respectively (Fig 6e,f). More detailed analysis of the electrophysiological data evidenced a significant decrease of total VT events/per animal in LV-Cx43 mice compared to the two control groups (Fig 6g). We also performed echocardiography and found that cardiac pump function was comparable in the different groups (Fig 6h,i), likewise infarct size (Suppl. Fig 5b,d). Interestingly, we could observe the impact of cardiac remodeling, as infarct areas were larger (Suppl. Fig 5b) and volumes smaller (Suppl Fig 5d) in 8 weeks, compared to 2 weeks hearts (Suppl. Fig 5a,c).

**Figure 6:**
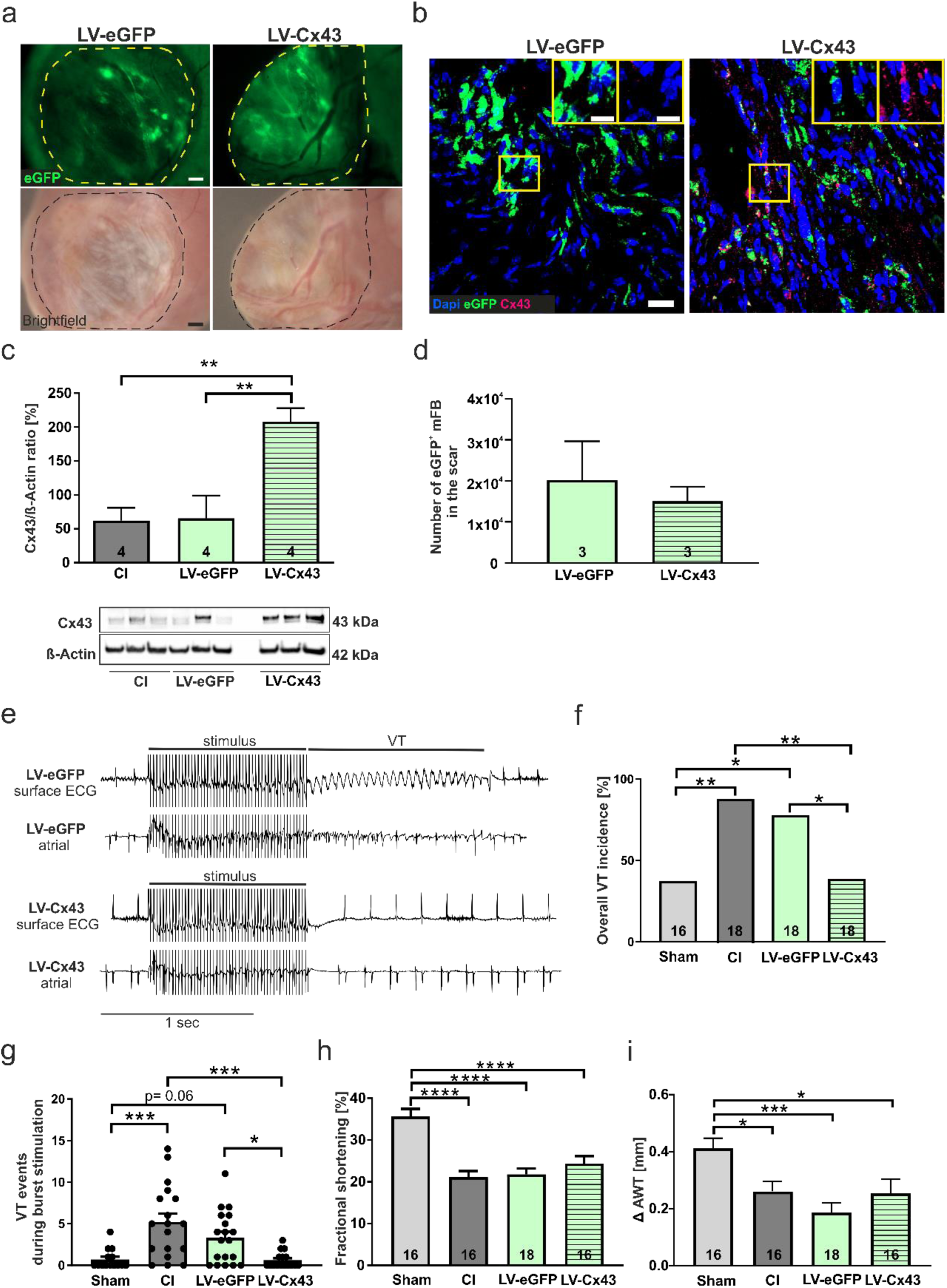
Long-term engraftment of mFB and functional impact at 8 weeks post-surgery. (a) Macroscopic pictures document engraftment of eGFP^+^ mFB in the scar area (dotted area, bars= 500 µm). (b) Anti-eGFP and Cx43 stainings of mFB in a LV-eGFP and a LV-Cx43 heart (blue= nuclei, green= eGFP, pink= Cx43, bars= 20 µm and 5 µm). (c) Cx43 protein expression in cut-out scars analysed by Western blotting. (d) Quantitation of engrafted eGFP^+^/Vimentin^+^ mFB. (e) Representative surface ECG traces and (f) VT incidence in the different groups of mice. (g) Analysis of VT events. h,i: Echocardiographic assessment of fractional shortening (h) and anterior wall thickening (i) in the different groups of mice. P-values: *= < 0.05; **= < 0.01; ***= < 0.001; ****= 0.0001.

Thus, our data demonstrate that grafting of Cx43 expressing mFB into cardiac infarcts strongly reduces short-and long-term electrical vulnerability *in vivo*.

## 5. Discussion

The cardiac scar has so far proven little amenable for targeting and/or gene therapy strategies. Herein we show an efficient approach to target and modulate the functional properties of the scar by combining *ex vivo* gene therapy in mFB with *in vivo* cell therapy with the help of magnetic steering. Fibroblasts account for 20-27 % of cardiac cells and are known to play an important biological role in the heart (28). Upon ischemic stress, many fibroblasts die, but surviving cells transdifferentiate into αSMA^+^ myofibroblasts, strongly proliferate and repopulate the cardiac scar representing the best surviving cell source within cardiac lesions (28, 29). Thus, cardiac myofibroblasts resist the adverse local conditions in ischemic lesions, stabilize these against rupture and play a key role in scar formation (28, 29). Given these distinct cellular properties of cardiac myofibroblasts and the so far futile efforts to efficiently target the cardiac scar, we probed a combination of gene and cell therapy using cardiac myofibroblasts. We opted for embryonic mFB given their strong proliferation rate enabling easy cell expansion *in vitro* for grafting experiments. Despite testing different AAV serotypes, in the past, we have not been able to transduce mFB with AAV, whereas roughly 40 % of cells were found eGFP^+^ upon LV transduction. Interestingly, in our earlier work we demonstrated that magnetic steering using MNP-loaded cells strongly enhanced eCM engraftment rates (6). We therefore used the same approach after testing different MNP. We identified PMAO-MNP as the best suited MNP, as they displayed a good magnetic moment, low cellular toxicity, no intracellular aggregate formation, and efficient degradation by lysozymes (15, 30).

Our initial experiments revealed at 2 weeks after injecting LV-treated 200.000 mFB into the fresh cardiac lesion a so far not observed degree of cell engraftment. Assuming that half of the injected cells are eGFP^+^, grafted mFB numbers would be close to 100.000, which are far more than those reported after injecting mesenchymal stem cells (31, 32) or cardiomyocytes (6, 33). Our data suggest that both, the tolerance of mFB towards the adverse conditions in cardiac lesions; and the lack of prominent cell death and the strong proliferation rate of these cells underlie these massive engraftment rates. In fact, almost 50 % of mFB were still proliferative after more than two weeks in cell culture and also a significant increase in Ki67^+^ cells were observed in the cardiac scar area after grafting mFB. Our experiments also showed that the MNP/magnet approach enhanced mFB engraftment rates 3-fold, underscoring the impact of this technology. Because less than 50 % of grafted cells were eGFP labelled, we took advantage of a double transgenic system to assess engraftment site and rates, the interaction of exo-with endoFB and the proliferation of the two cardiac myofibroblast subpopulations. For these experiments neonatal cardiac myofibroblasts (exoFB) were used. Because of the low recombination rate upon Tamoxifen administration, for most of the experiments tomato fluorescence was used as genetic marker. Analysis of the exoFB at different time points of cell culture revealed that the *in vitro* enrichment yielded >95 % cardiac myofibroblasts and that very few tomato^+^ endothelial cells and no cardiomyocytes were found after grafting. Numbers of grafted cells were confirmed to be very high at 1 and 2 weeks post-surgery, but lower than the extrapolated numbers for mFB, which is consistent with the approximately 4-fold lower *in vitro* proliferation rate of exoFB compared to mFB. We found that the exoFB were preferentially located in the epicardial area of the lesion, whereas the endoFB were positioned close to the endocardial layer, this is probably due to the intracardiac injection method. Surprisingly, endo-and exoFB did not migrate and mix over time, but rather stayed closely aligned in the above described location. Importantly, grafting of exoFB enhanced the proliferation rate of endoFB by 4-fold suggesting signaling between these two cell populations despite apparent lack of direct cellular interactions. Given the prominent engraftment rates of mFB in the cardiac lesion we interrogated, whether a combination of *ex vivo* gene and *in vivo* cell therapy modulates the functional properties of the infarcted heart. We have shown in our earlier work that grafting of Cx43^+^ excitable cells (7) or direct LV-mediated transduction of resident endoFB (3) strongly reduced post-infarct VT incidence *in vivo*, even though direct LV transduction was very poor. This anti-VT protective mechanism is due to a modest increase of conduction velocity in the border zone and the scar area, which is sufficient to strongly prevent reentry events resulting in VT at least in the small scars of the mouse heart. The LV-mediated Cx43 overexpression of mFB was found in cell culture and after grafting of mFB at gene and protein levels. Interestingly, Cx43 overexpression in mFB led to a partial myogenic transdifferentiation of the cells based on the gene expression pattern, but not at the protein level. The grafting of Cx43 overexpressing mFB resulted in a prominent reduction of VT incidence at 2-and 8 weeks post-surgery, despite the clearly lower number of grafted mFB at the late time point. The causes responsible for the loss of grafted cells over time is unclear, we suspect that immune-based cell rejection could be an important factor, as immunosuppression was stopped at 10 days post-surgery. In addition, we also suspect normal cell loss, as the cardiac scar is thinning out (see also Suppl. Fig 5), becoming less cellularised over time. In fact, histological and functional data showed increased cellular content and left anterior wall thickening at 2 weeks after cell injection resulting in improved left ventricular function, probably because of changes in wall stiffness, whereas this was not anymore observed at 8 weeks post-surgery.

In summary, we present a novel approach of combined *ex vivo* gene and *in vivo* cell therapy with mFB in combination with MNP/magnetic steering to efficiently target the cardiac scar and modulate the function of the heart. The efficacy of this approach is due to the resistant nature of cardiac myofibroblasts and their proliferation characteristics after grafting *in vivo*. Engraftment was also strongly increased because of PMAO-MNP in combination with magnetic steering. Cardiac myofibroblasts have been reported to have beneficial effects on the cardiomyogenic potential of pluripotent stem cells, the grafting of Embryonic-Stem (ES)-cell-derived CM, scar formation and its mechanical stabilization (34). The observed engraftment rates are much higher (∼4-fold) compared with other cell types such as eCM. Of further advantage is that autologous cardiac myofibroblasts could be obtained using biopsy outgrowths or the *in vitro* differentiation of patient-specific hiPSC, while we have not observed any adverse effects such as migration and increased fibrosis outside of the scar. This strategy could be, for instance, used to modify the immune response in the scar by overexpressing immunomodulating factors or other cell biological and/or functional targets in mFB. Future work should explore strategies to maintain engraftment rates more stably, either by using autologous cells and/or expressing antiapoptotic genes. Moreover, it would be important to assess, whether embryonic or postnatal myofibroblasts are best suited and whether long term engraftment and improved functional effects can be also achieved in large animal models.

## Author Contributions

M.S. isolated/transduced mFB, performed surgeries, electrophysiological testing, Western blots, immunostainings, and analysis of data. K.W. tested the loading of mFB with various MNP *in vitro*. E.C. also performed surgeries and electrophysiological testing, J.N. and M.M. performed *in vitro* characterisation and analysis of double-transgenic neonatal exoFB. M.H. analysed the bulk RNA-seq Data, R.F. and J.M.D.F. provided and characterised different MNP. S.H. and A.P. generated LV constructs and provided LV for *in vitro* and *in vivo* experiments, D.E. performed magnetic particle spectrometry. B.K.F. and W.R. designed the study, analysed data and wrote the manuscript.

All authors have read and agreed to the published version of the manuscript. B.K.F. and W.R. have equally contributed to this work.

Funding: This study was part of SFB 1425, funded by the Deutsche Forschungsgemeinschaft (DFG, German Research Foundation)—project # 422681845 (W.R. and B.K.F.).

Institutional Review Board Statement: Not applicable.

Informed Consent Statement: Not applicable.

Data Availability Statement: Not applicable.

## Acknowledgments

We thank the members of the Institute of Physiology I (Bonn Germany), namely P. Freitag and A. Markert for excellent technical assistance, T. Mohr and P. Niemann for support in histological analyses, C. Geisen and S. Rieck for technical input on LV titration and LV/MNP complex formation. We thank M. del Puerto Morales (Madrid, Spain), C. Plank and O. Mykhaylyk (Munich, Germany) for providing different types of MNP. We also thank the Flowcytometry Core Facility of the Medical Faculty of the University of Bonn for support.

Conflicts of Interest: The authors declare no competing interest.

## Notes

### Competing Interest Statement

The authors have declared no competing interest.

