## Supplemental Figures for "Efficient targeting of heart lesions with cardiac myofibroblasts: Combined gene and cell therapy enhanced by magnetic steering"

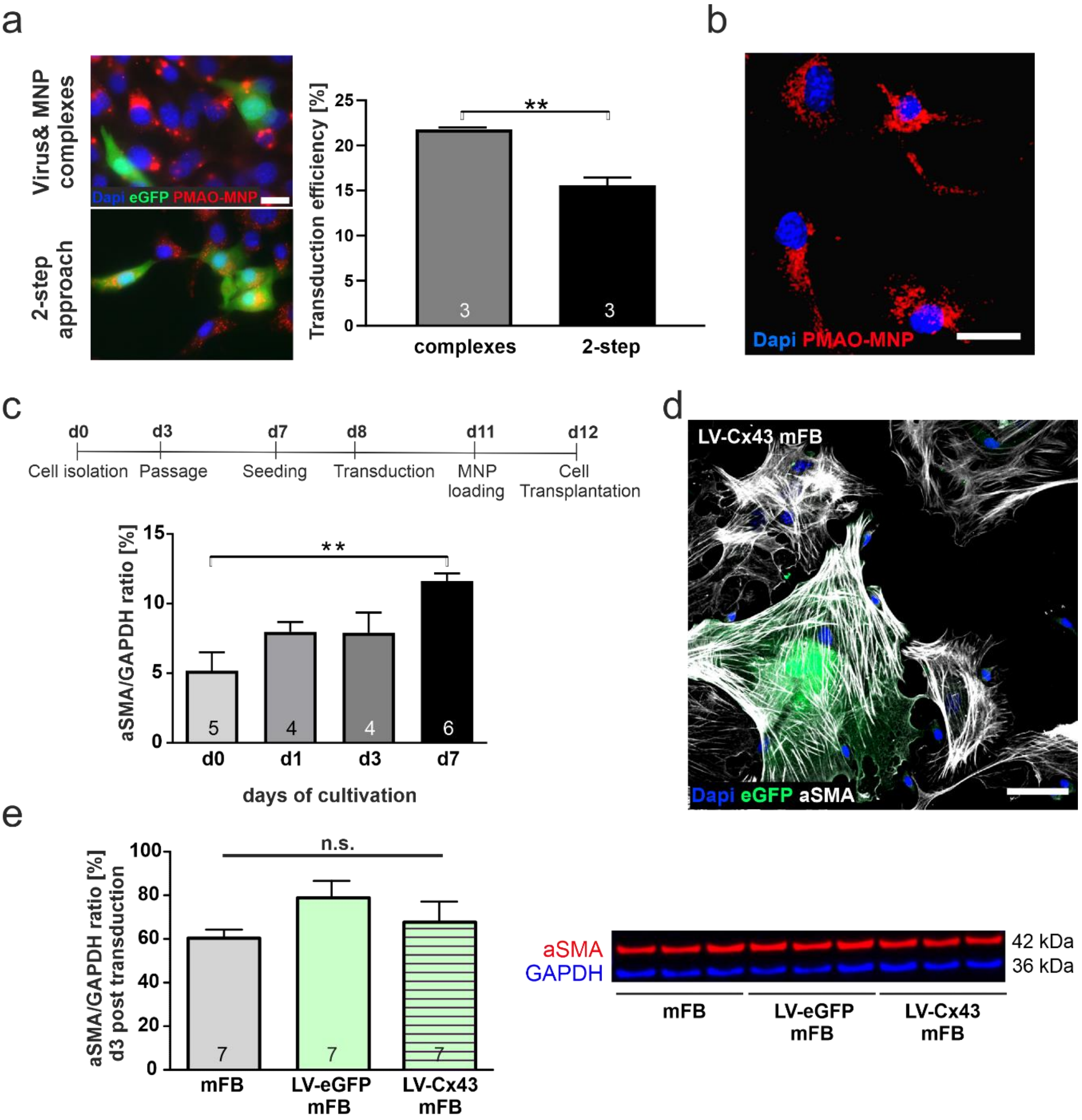


**Supplementary Figure 1: Lentiviral (LV) transduction and MNP loading of Fibroblasts *in vitro*.** (a) Combined LV-transduction and MNP loading of 3T3-FB with PMAO-MNP/ LV complexes (left picture, 30 min, at 37 °C in HBSS$^{++}$, + magnet) or a 2-step approach with initial LV transduction, followed by PMAO-MNP loading after 48 h (right picture, overnight, at 37 °C in cell culture medium, - magnet) (blue= nuclei, green= eGFP, red= PMAO-MNP, bar= 20 µm). (b) Murine embryonic cardiac fibroblasts (mFB) following PMAO-MNP loading (25 pg Fe/cell (blue= nuclei, red= PMAO-MNP, bar= 20 µm)). (c) Time-dependent αSMA protein expression of freshly isolated mFB during cultivation (analysed by Western blotting). (d) Histological image of LV-Cx43 mFB at d3 post transduction (bue= nuclei, green= eGFP, white= αSMA, bar= 50 µm). (e) αSMA protein expression of LV-treated and control mFB (analysed by Western blotting) at 3 days post transduction. P-value: **= < 0.01.

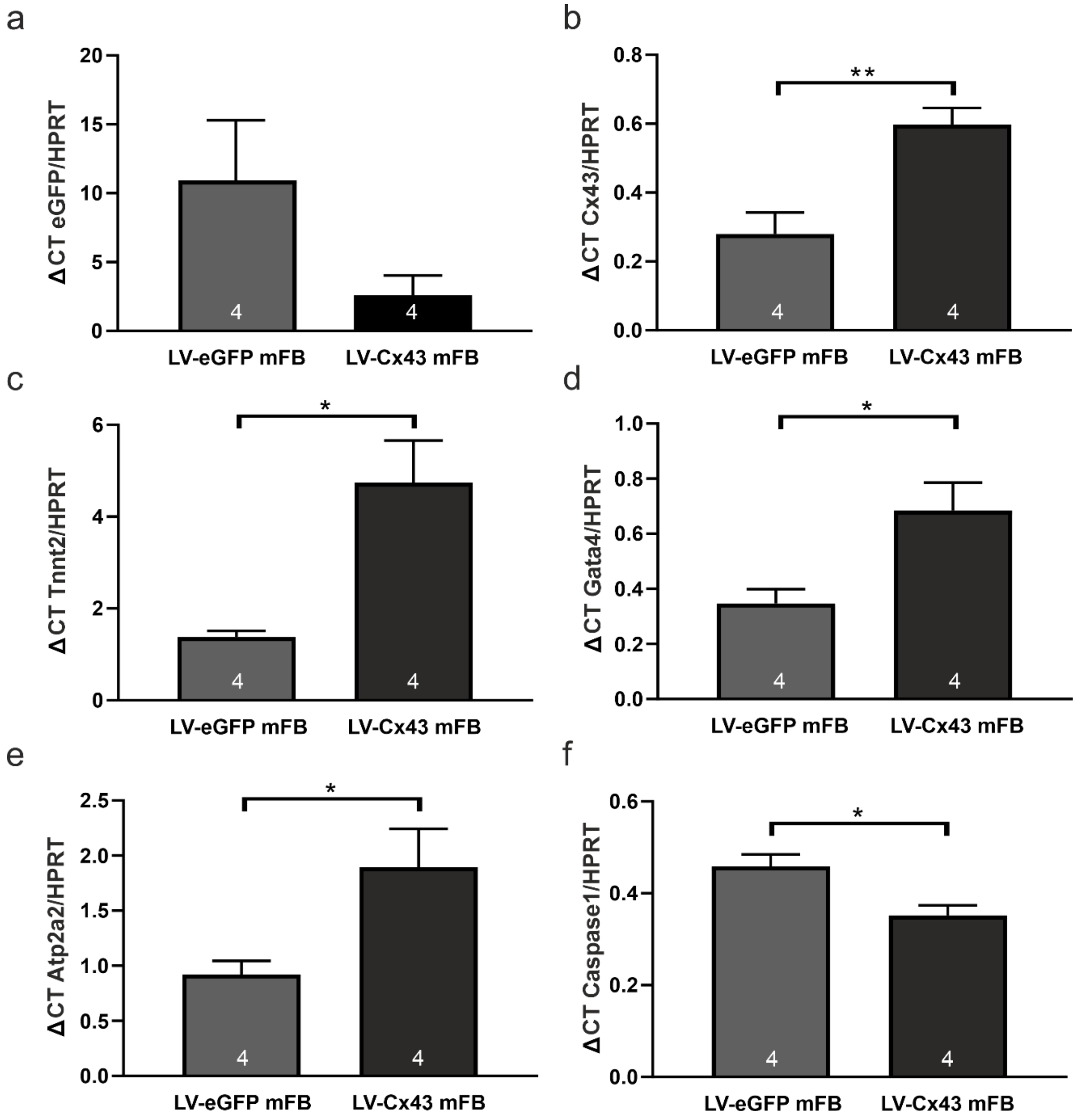


**Supplementary Figure 2: qPCR of LV-eGFP and LV-Cx43 transduced mFB.** LV-eGFP and LV-Cx43 (MOI=5, 25 pg Fe/cell PMAO-MNP) at 3 days post transduction. (a) eGFP, (b) Cx43, (c) Tnnt2, (d) Gata4, (e) Atp2a2 and (f) Caspase1 assays. Graphs show ΔCTvalues in relation to the housekeeping gene HPRT. P-values: *= < 0.05; **= < 0.01

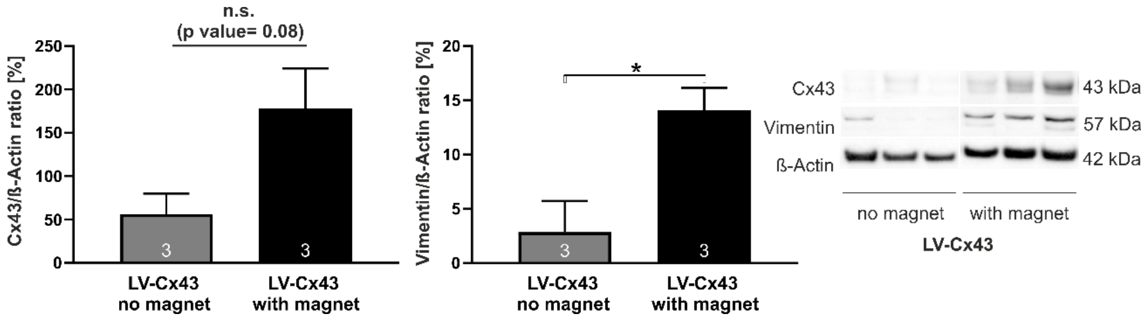


**Supplementary Figure 3: Enhanced cell engraftment assessed by Western Blotting.** (a) Cx43 and Vimentin protein content in excised scar tissue of LV-Cx43 hearts with and without magnet application at 2 weeks post-surgery. P-value: *= < 0.05

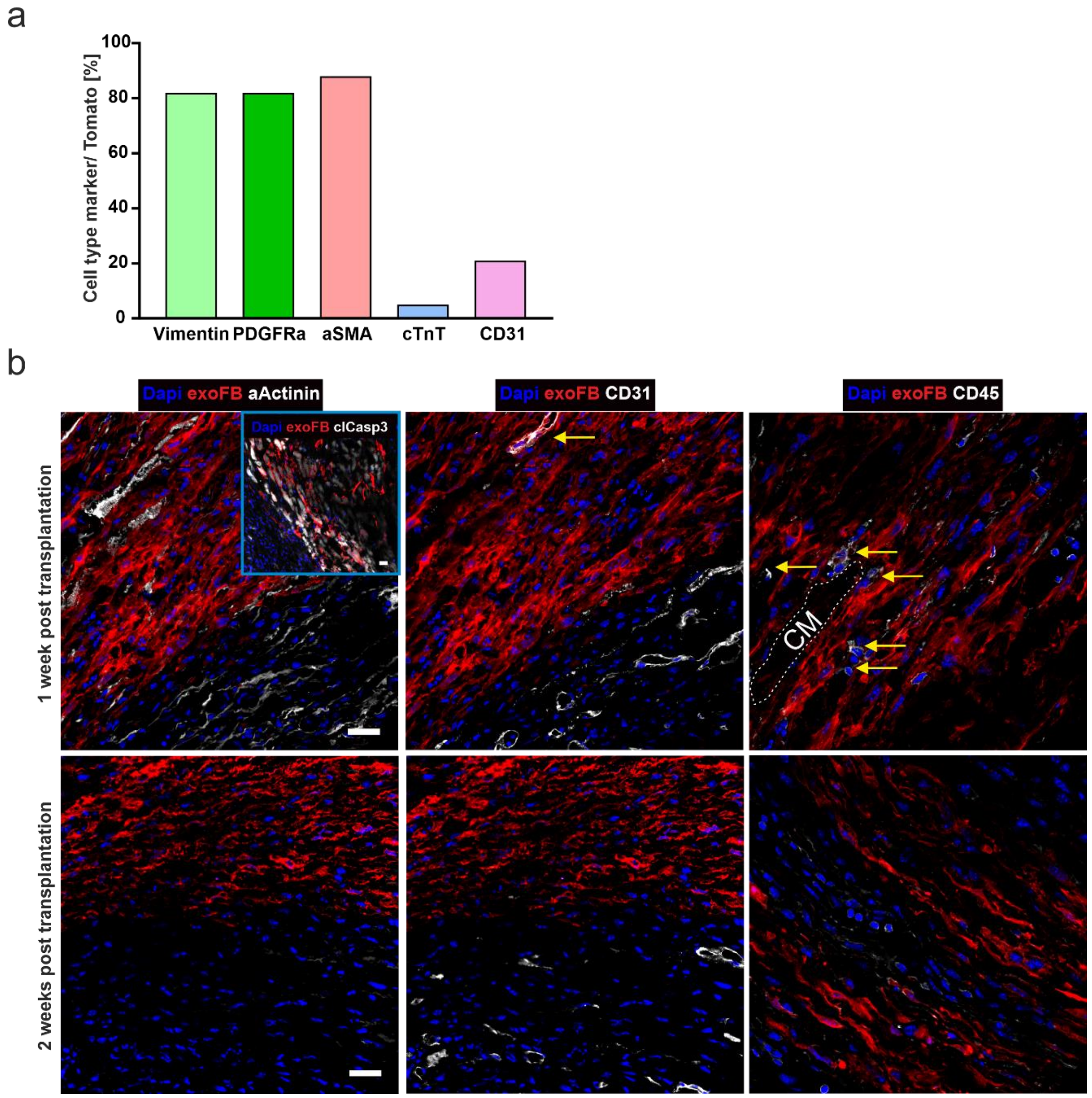


**Supplementary Figure 4: Cellular characteristics of the myocardial lesion at 1 and 2 weeks following injection of exoFB obtained from double-transgenic mice.** (a) Immunohistochemically characterisation of cells harvested from Tamoxifen-induced double-transgenic mTmGxTcf21$^{MCM}$ hearts after 5 days of cell culture. (b) Few tomato$^{-}$/$\alpha$actinin$^{+}$ apoptotic (clCasp3$^{+}$) cardiomyocytes, tomato$^{+}$/CD31$^{+}$ endothelial and tomato$^{-}$/CD45$^{+}$ immune cells are detected at 1 week after exoFB injection (upper panels). At two weeks post-injection, no clCasp3$^{+}$, no tomato$^{+}$/CD31$^{+}$ and only very few tomato$^{-}$/CD45$^{+}$ cells are observed (lower panels). bars= 20 µm.

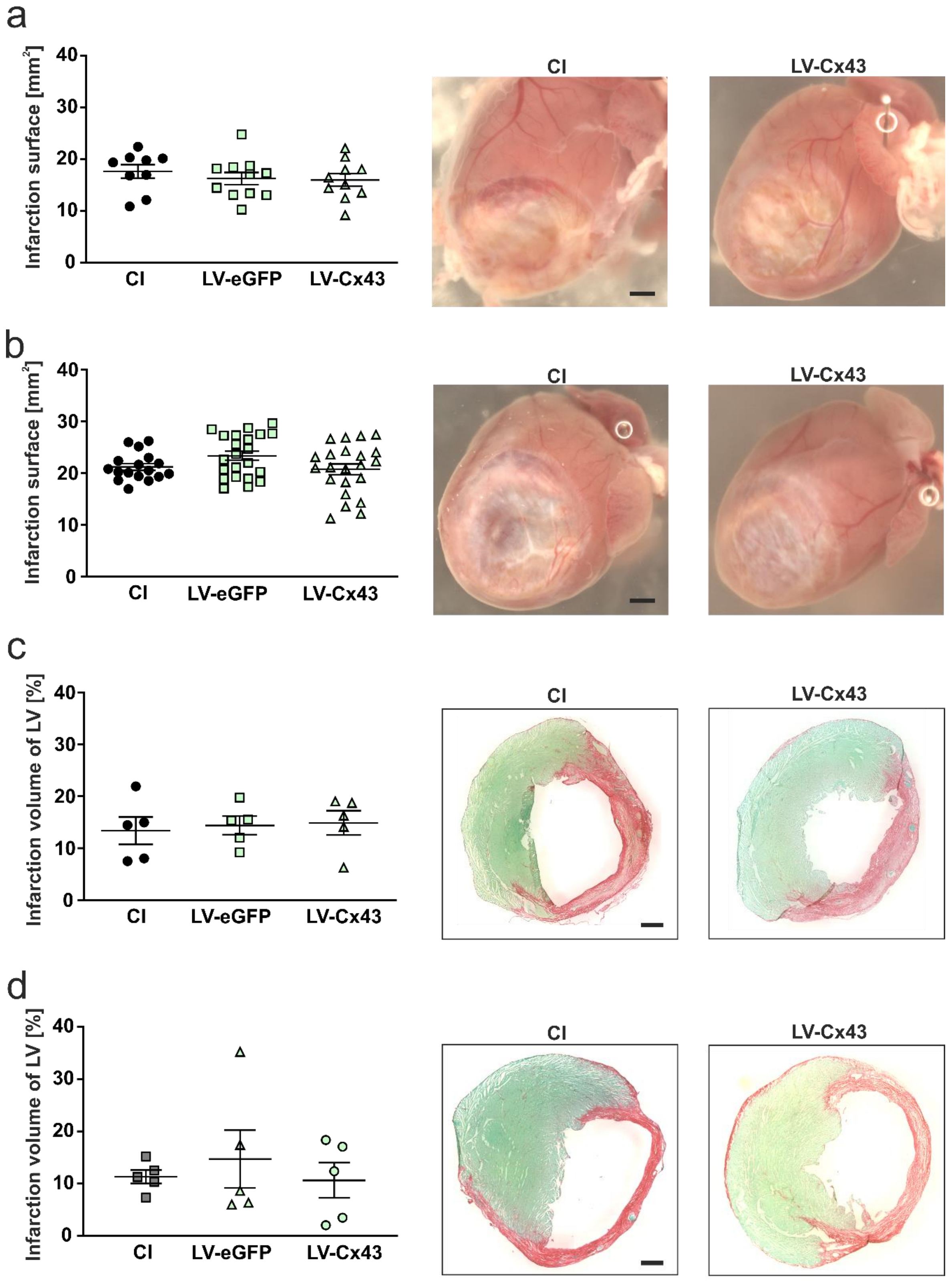


**Supplementary Figure 5: Characterisation of the myocardial lesion (CI) at 2- and 8 weeks after injecting LV-transduced mFB**. Analysis of infarction surface at (a) 2 and (b) 8 weeks post-surgery. Analysis of infarction volume at (c) 2- and (d) 8 weeks post-surgery. a, b: bars= 1000 µm; c, d: bars= 500 µm.
